# Lossless and Contamination-Free Digital PCR

**DOI:** 10.1101/739243

**Authors:** Peiyu Liao, Mengcheng Jiang, Zitian Chen, Fangli Zhang, Yue Sun, Jun Nie, Meijie Du, Jianbin Wang, Peng Fei, Yanyi Huang

## Abstract

The realization of the vast potential of digital PCR (dPCR), to provide extremely accurate and sensitive measurements in the clinical setting, has thus far been hindered by challenges such as assay robustness and high costs. Existing popular dPCR platforms that target the clinic have not reached wide-spread adoption, due to problems with sample loss and risk of contamination during sample preparation, compartmentalization, and transfers; limitations of dynamic range and signal-to-noise in the result readout also restricts broad applications. Here we introduce a lossless and contamination-free dPCR technology termed CLEAR-dPCR, which addresses these challenges by completing the dPCR sample preparation, PCR reaction, and readout all in one tube. We achieve this by adjusting the refractive index of the aqueous PCR mix to make the emulsion optically transparent, and devised a light-sheet microscope to capture 3D images of the cleared emulsion for results readout. This approach demonstrates improved accuracy over existing dPCR platforms, and enables a greatly increased dynamic range to be comparable to that of real-time quantitative PCR (qPCR). CLEAR-dPCR is an easy to operate, sensitive and accurate dPCR platform that we envision will fulfill the potential of dPCR for routine use clinical diagnosis.

## Main Text

The desire for better quantitative assessment of genetic information is always increasing. Techniques for sequence-specific nucleic acid quantification are key tools in biology and medicine, especially in scenarios involving rare genetic variant detection where both high sensitivity and high specificity are warranted ^1-3^. Unlike real-time quantitative PCR (qPCR), where relative quantification is achieved using a standard curve, digital PCR (dPCR) ^4^ enables absolute quantification with up to single-molecule resolution. Soon after the first demonstration of dPCR through microtiter plate compartmentalization ^4,5^, various improvements using microfluidics-based methods were developed ^6-10^. Nanoliter or sub-nanoliter ^11^ microfluidic compartmentalization reduces reagent cost, increases reaction efficiency, facilitates large dynamic-range Poisson distribution, and simplifies the experimental process ^12-15^.

A typical dPCR process comprises three major steps: partitioning, amplification, and counting. Partitioning can be implemented via droplets ^9,16^ or micro-chambers^12^. Counting is also done via two major approaches: serial reading and planar imaging. Serial reading naturally fits the transferability of the fluidic emulsion and has been widely coupled with droplet-based compartmentalization. Meanwhile, planar imaging is more compatible with array-based compartmentalization ^13^.

Currently, droplet dPCR is gaining in popularity ^14^, partly because the emulsion can be easily transferred into conventional microcentrifuge tubes and the amplification can be conducted in conventional PCR machines. However, the prevailing flow-focusing-based microfluidic droplet generation strategy often used in this scenario suffers from sample loss during the flow stabilization period. When serial counting is applied, further sample loss and contamination are inevitable during sample transfer from the PCR tube to the reading device. Recently we have demonstrated the use of micro-capillary array (MiCA) for lossless emulsion generation (Supplementary Fig. 1) and subsequent dPCR ^15^. However, the counting step employed a serial counting strategy that still suffered from the same sample loss and contamination issues as other dPCR strategies. Here, we demonstrate a novel dPCR strategy, characterized by centrifugation-based droplet generation, optical clearing of PCR emulsion, and *in situ* counting based on light-sheet fluorescence microscopy (Figure 1). With this novel approach, named CLEAR-dPCR, we are able to achieve lossless and contamination-free dPCR.

**Fig. 1.**
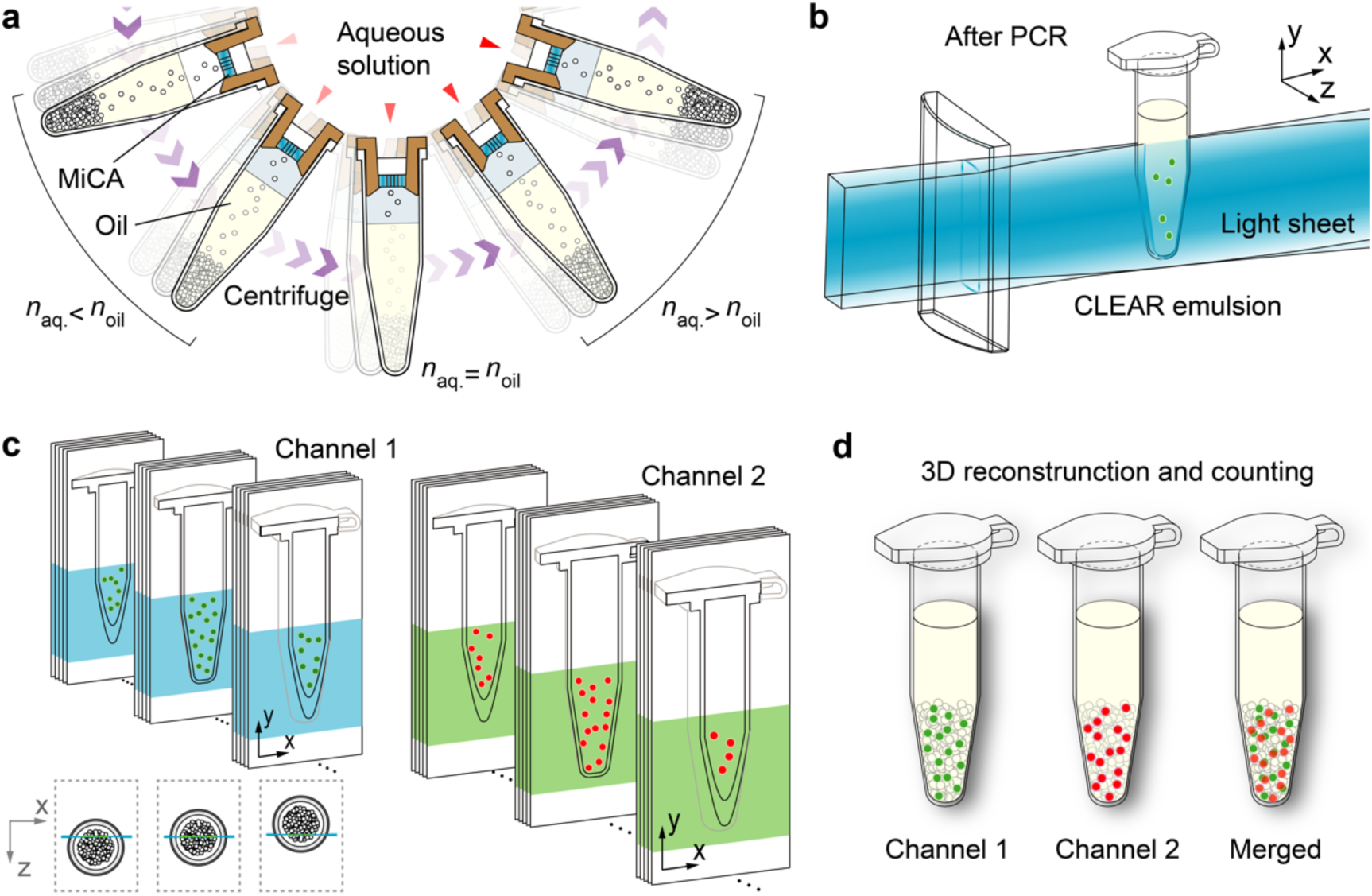
Schematic illustration of CLEAR-dPCR process. **a**, The generation of droplet emulsion using MiCA. Droplet PCR samples are centrifuged in a swing-bucket rotor and jet through the through-holes on MiCA plates, yielding massive droplets monodispersed in silicone oil. CLEAR emulsion is formed by matching aqueous and oil phases’ refractive indices. **b**, The high-throughput readout of bulk PCR droplets, optically transparent and densely packed, in the PCR tube using light-sheet illumination. A laser light-sheet scans the droplets in the PCR tube *in situ*, and the plane fluorescence images are sequentially acquired using a high-speed sCMOS camera. **c**, The dual-channel light-sheet fluorescence image sequences, in which all the fluorescently positive droplets are recorded (red: HEX/VIC, green: FAM). **d**, The volumetric reconstruction of dual-channel images, and accurate droplet counting can be implemented based on the 3-D data.

## Results

### CLEAR digital PCR

In order to achieve lossless counting via optical readout *in situ*, we generate an optically transparent dPCR emulsion termed CLEAR emulsion, by dispersing aqueous dPCR mix with a refractive index close to that of the oil phase using MiCA centrifugation (Fig. 1a). After PCR, *in situ* noninvasive and high-throughput readout of whole-tube emulsion is accomplished by rapid 3D light-sheet fluorescence imaging (Fig. 1b) of the same tube. By recording a series of planar images of the CLEAR emulsion (Fig. 1c), we are able to rapidly reconstruct the 3D structure of the whole PCR emulsion and achieve digital counting of the positive droplets (Fig. 1d). This high-throughput approach streamlines dPCR with a convenient operation, thoroughly eliminates sample loss and contamination, and greatly reduces the cost of dPCR assays.

### Transparent emulsion via refractive index matching

In typical water-in-oil (w/o) emulsions, the massive number of droplets densely packed in the PCR tube creates an opaque emulsion due to the light scattering at the water-oil interfaces. We serendipitously discovered that a common PCR additive, betaine ^16^, effectively increases the refractive index of aqueous solution to be similar to that of the carrier oil, thereby minimizing the light scattering in the PCR emulsion. We carefully titrated the betaine addition from 0 to 5 mol/L (Fig. 2a, Supplementary Fig. 2) and found that an optimized concentration of 3.14 mol/L generates a PCR mix with a refractive index similar to that of the oil-surfactant mix (low-viscosity silicone, *n* = 1.3903). At this betaine concentration, the whole emulsion is completely transparent (Fig. 2b). Bright-field photographs show the highest light transmission rate (∼85%) at 3.14 mol/L. As a result of the greatly improved clarity, this optimized CLEAR-dPCR emulsion exhibited the highest image quality and signal-to-background ratio in light-sheet fluorescence imaging (Fig. 2c). We further verified that PCR reactions inside these CLEARed picoliter droplets are not affected by the addition of betaine (Supplementary Fig. 3).

**Fig. 2.**
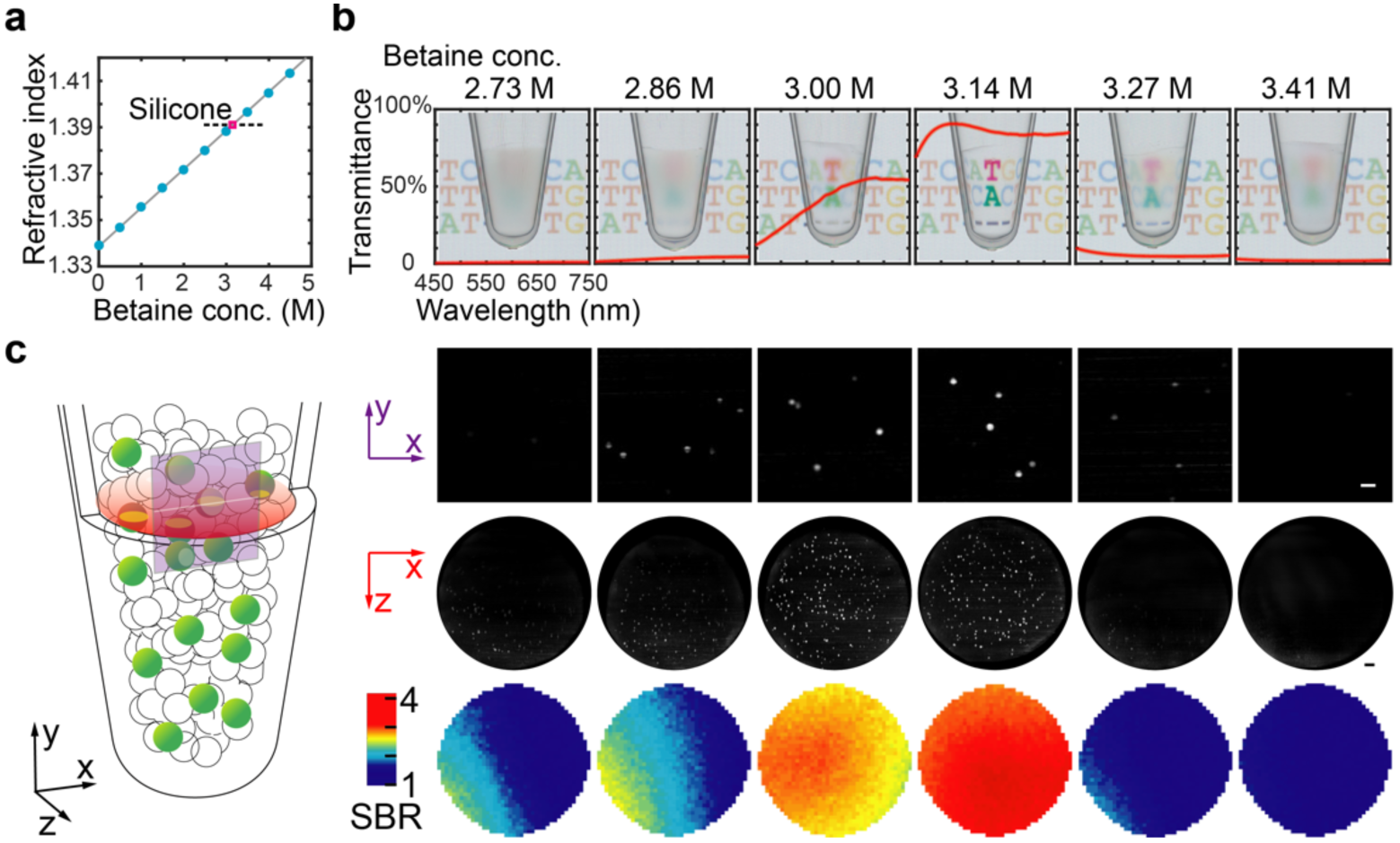
The optical clearing of the dPCR emulsion, which substantially improves the transparency and enables large-depth, whole-tube scanning. **a**, The change of PCR mix’s refractive index versus the concentration of betaine. **b**, Bright-field transmission images of the emulsions generated with different concentrations of betaine in PCR reaction buffer. The red lines plot the emulsions’ transmission spectra (light path: 1 cm). 3.14 mol/L betaine provides the best refractive index matching. **c**, The fluorescence images of emulsions at different betaine concentrations. Top: light-sheet fluorescence images taken by camera. Middle: reconstructed x-z plane images of the emulsions. Bottom: signal-to-background ratio distribution in x-z plane. Scale bars: 200 µm.

### Light-sheet sectioning the transparent emulsion

We built a compact light-sheet illumination microscope (Fig. 3a,b) to read the fluorescence signals in the bulk CLEAR-dPCR emulsion layer by layer. Our light-sheet readout geometry was customized for PCR tube scanning (Supplementary Fig.4). Dual-wavelength (488/532 nm) 3D-imaging was implemented for duplex dPCR with FAM and HEX/VIC labelling. The combination of plane illumination and a sCMOS camera provided a large imaging field of view of ∼4.6×3.6 mm, covering the entire emulsion (Fig. 3c). Together with a rapid acquisition rate of 100 frames/second, high-speed scanning of the whole tube can be completed within 6 seconds. The spatial resolution of our light-sheet imaging is 6.5 µm (lateral) and 20 µm (axial); sufficient to identify the 41-µm droplets in all three dimensions (Supplementary Fig. 5). By stacking the raw light-sheet images into a 3D volume, we further identified and counted the positive droplets using a customized algorithm (Fig. 3d and Supplementary Fig. 6).

**Fig. 3.**
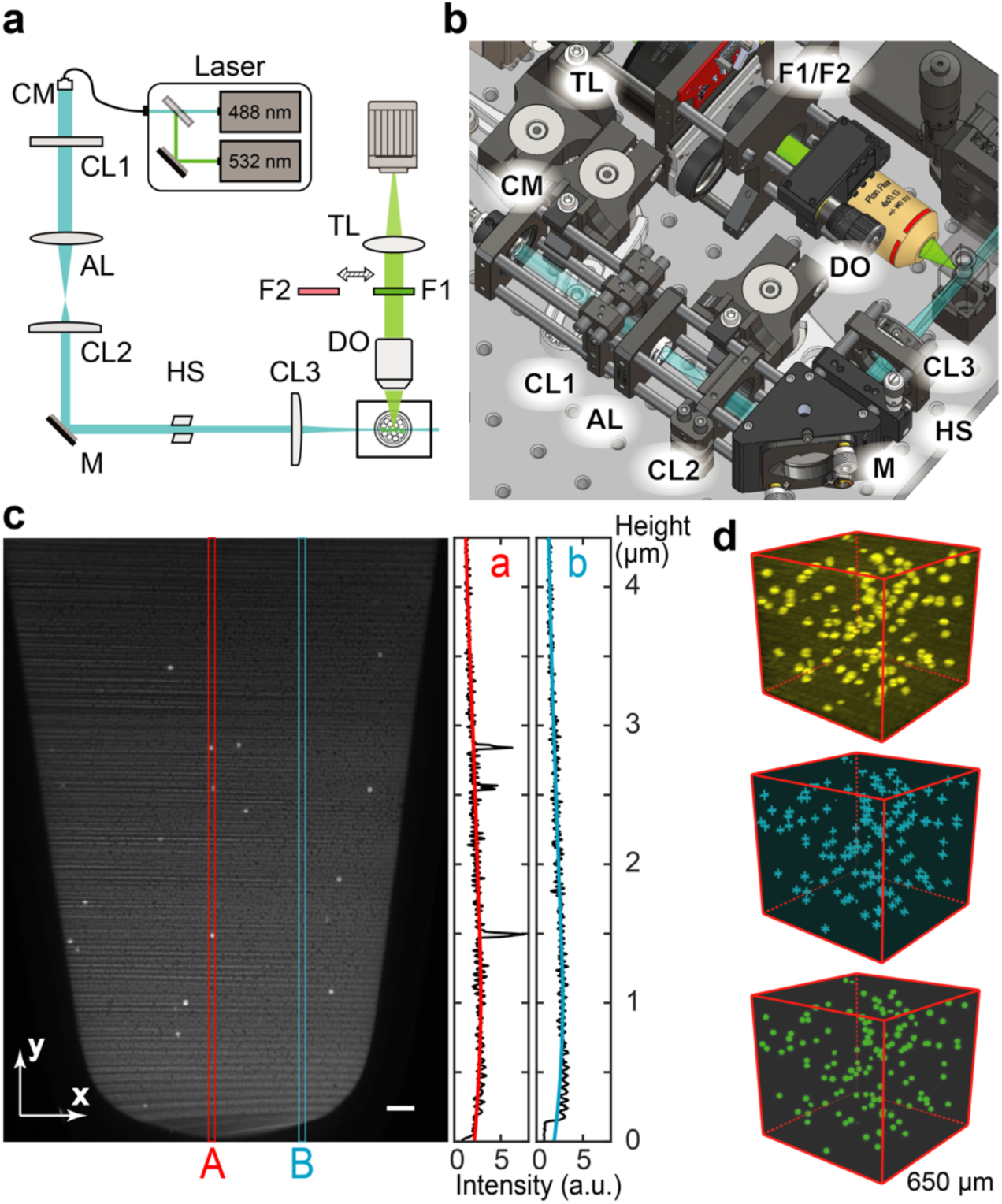
The light-sheet fluorescence reader and droplet counting. **a**. The schematic light path and major components. A wide laser light-sheet which can illuminate the entire emulsion in PCR tube is generated by the line focusing of a collimated elliptical beam. CM: collimator, CL: cylindrical lens, AL: aspherical lens, HS: horizontal slit, DO: detection objective, F: filter, TL: tube lens. **b**. The compact (30×30 cm) layout of the setup. **c**. Signal and background measurement in a raw image of CLEAR-dPCR. The red line cut **A** includes three positive droplets while the blue line cut **B** includes only the background. Uneven light field will be further corrected and the background signal will be subtracted. Scale bar: 200 µm. **d**. The 3D reconstruction of dPCR images. Raw images are first reconstructed into a 3D intensity matrix (top, yellow). Then *in silico* scanning detects the spatial local intensity maxima (middle, cyan crosses) as candidate positive signals. Finally, the fluorescently positive droplets (bottom, green dots) are located and counted based on the identified candidates.

To evaluate the performance of CLEAR-dPCR, we used three assays to benchmark it against dPCR industry leaders: Bio-Rad droplet dPCR (QX200-ddPCR) and ABI 7500 Fast Real-Time PCR (7500F-qPCR).

### Absolute quantification

We performed absolute quantification of a 280-bp amplicon DNA from *Listeria monocytogenes* genome (Supplementary Table 1). We serially diluted the template DNA to different concentrations spanning five orders of magnitude (10^1^–10^5^ copies/aliquot). The results show that CLEAR-dPCR and QX200-ddPCR both had excellent linearity (R=0.9999) within the range of 10-1.3×10^5^ copies/aliquot, while 7500F-qPCR demonstrated lower linearity (R=0.9975) (Fig. 4a,b and Supplementary Table 2-4). 7500F-qPCR quantitation relies on a standard curve and some of the measurements apparently deviated from those obtained from the digital methods with much lower precision (Fig. 4c and Supplementary Table 4), which makes 7500F-qPCR unsuitable for absolute counting. The relatively higher measurement uncertainty of low copy number samples was mainly due to sampling error, which is independent of specific methods.

**Fig. 4.**
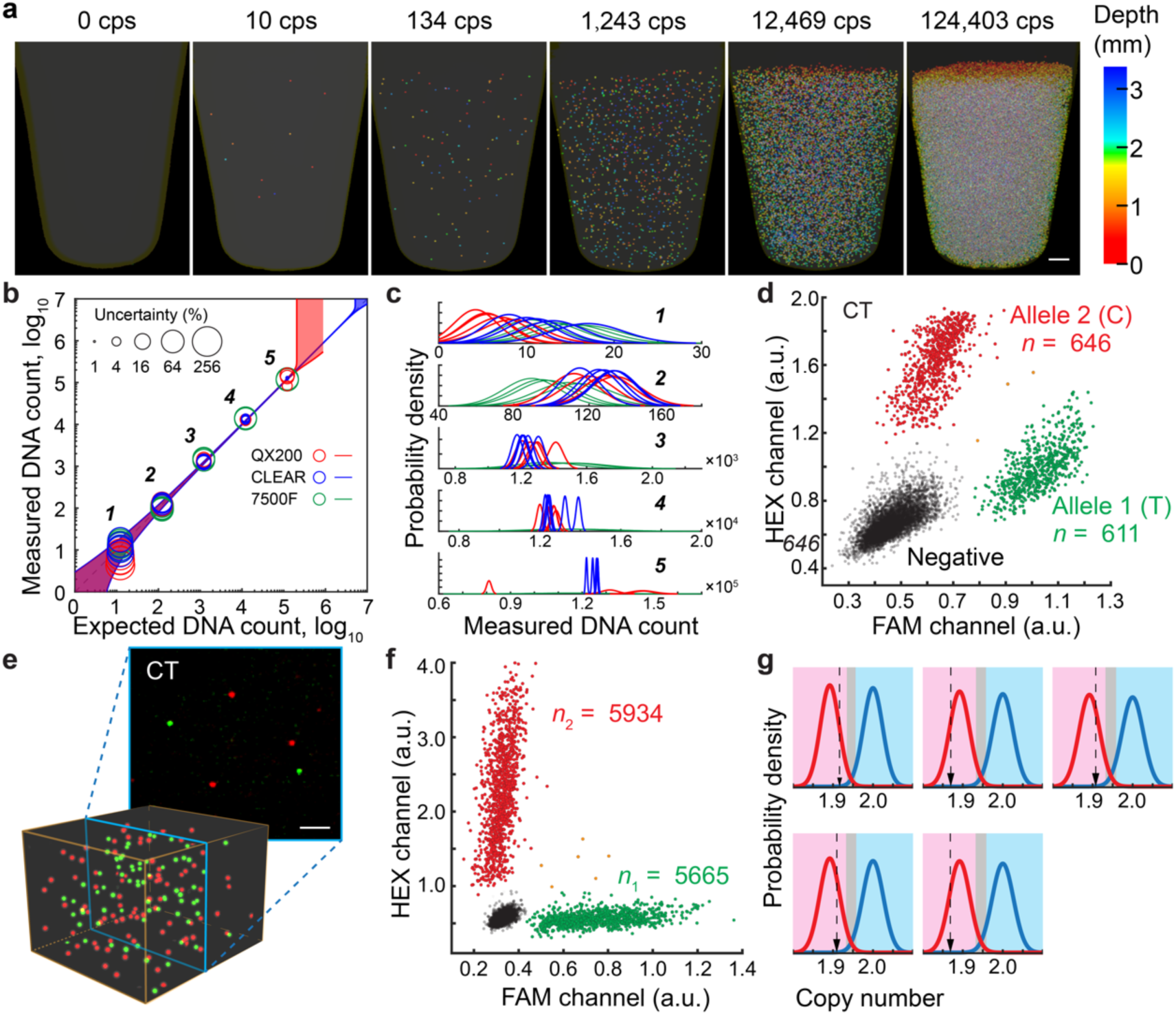
Performance of CLEAR-dPCR with different assays. **a**, Absolute *in situ* quantification of positive droplets in PCR tube, with 10^5^ dynamic range. Scale bar: 500 µm. **b**, Comparison of quantification performance by CLEAR-dPCR, QX200-ddPCR and 7500F-qPCR, with their uncertainty levels (95% confidence) shown as the circle sizes. The shaded areas along the diagonal represent the 95% confidence boundaries. **c**, Probability density distribution of each measurement. **d**, SNP detection and allele counting. Scatterplot results of a heterozygous sample at SNP rs10092491. **e**, The heterozygote’s merged images (top) and 3D reconstructions (bottom) of both channels. Scale bar: 200 µm. **f**, Clinical copy number variation detection. Copy numbers of Chr.1 uc13 (red) and *TSC2* gene (green) were detected in a mock sample with 5% deletion of *TSC2* exons. **g**, Results of 5 TSC tests of mock affected samples using a likelihood ratio classifier. Relative copy number in the magenta region is classified as affected, and in light blue as healthy, otherwise in grey undetermined.

It is worth noting that, while both dPCR systems presented equivalent linearity, QX200-ddPCR measurements had higher uncertainty for the more concentrated samples (Fig. 4b,c, 10^4^ and 10^5^ copies/aliquot). We attributed this to the combination of low droplet number and high sample loss of QX200-ddPCR. As shown in Supplementary Figure 7, target molecule distribution in QX200-ddPCR becomes over-saturated when the expected copy number approaches 10^5^; in this regime, the standard Poisson correction is insufficient and leads to error. With nearly half a million droplets and complete elimination of sample loss, CLEAR-dPCR maintains reasonable uncertainty with over 10^6^ target molecules (Fig. 4b). This dynamic range is comparable to that of qPCR (*Ct* value 15-33) and typically not feasible in dPCR; our CLEAR-dPCR method is distinctly advantageous in maintaining absolute quantification while greatly expanding the dynamic range of dPCR.

Precise measurement of low copy number targets further relies on low false positive rates. In the second benchmark assay, we investigate possible false-positive counting against a high background, by using human genomic DNA as template instead of *Listeria* DNA in same way as the first assay. When approximately 60 fg of DNA (equivalent to 20 copies of the human genome) per droplet was used as the input, we found no false-positive signals in the CLEAR-dPCR results.

### Single nucleotide resolution

Digital PCR has been intensively applied to clinical diagnostics for its unrivaled accuracy, which is an essential requirement to identifying single nucleotide variation in clinical samples, such as in circulating DNA-based liquid biopsy ^1,2,17,18^. For the third benchmark, we designed a TaqMan genotyping assay with two probes, each targeting an allele of SNP rs10092491 with FAM or HEX labeling (Supplementary Table 5 and Supplementary Fig. 8). Genomic DNA from eight human peripheral blood samples was quantified by two-color CLEAR-dPCR and QX200-ddPCR, yielding three types of positive droplet patterns for the three different genotypes (Figure 4d, Supplementary Fig. 9 and Supplementary Table 6). For heterozygous samples, both channels yielded almost equal numbers of positive counts. CLEAR-dPCR demonstrated higher accuracy with a mean count ratio of 99.6%, compared with the ratio of 104.9% for QX200-ddPCR. By checking the two-color 3D images taken from the heterozygous samples, we discovered that most positive signals in the two channels did not overlap (Fig 4e and Supplementary Fig. 10). A few droplets were positive in both channels, which is consistent with the Poisson statistical expectation (Fig. 4d). For homozygotes, both CLEAR-dPCR and QX200-ddPCR detected zero false positive signals (Supplementary Table 6).

### Application in copy number quantification

We further evaluated CLEAR-dPCR on copy number variation (CNV) measurements. The detection relies on measurement of the relative abundances of a target region and a reference region. We first tested the performance of CLEAR-dPCR duplex digital counting on sex determination by Y chromosome copy number quantification. Two TaqMan assays targeting the *SRY* gene on Chr Y (FAM) or ultra-conserved region (uc) 13 on Chr 1 (HEX) were used (Supplementary Table 7). Both CLEAR-dPCR and QX200-ddPCR confirmed the presence of Chr Y in all male samples, with count ratio between Chr Y and Chr 1 approaching 1:2 (Supplementary Fig. 11 and Supplementary Table 8). CLEAR-dPCR demonstrated comparable accuracy to QX200-ddPCR. In female samples, CLEAR-dPCR detected zero Chr Y signals among 3,074 Chr 1 signals, while QX200 ddPCR gave one false positive Chr Y count among 1,341 Chr 1 signals (Supplementary Table 6).

### Prenatal fetal DNA exonal deletion identification

Prenatal genetic diagnosis using cell-free DNA in maternal blood is an area where dPCR can play an indispensable role. Existing noninvasive prenatal test methods only focus on large CNV or point mutations. Exon deletion has rarely been covered due to technical difficulties ^19^, despite its essential role in many diseases ^20,21^. With CLEAR-dPCR, we demonstrated an exon copy number counting assay for tuberous sclerosis complex (TSC), an autosomal dominant Mendelian disorder caused by mutations in the *TSC1* or *TSC2* gene. Owing to the relatively large sizes of the two genes, exonal deletions are present in as many as 10% of clinical cases ^21^. A previous multiplex ligation-dependent probe amplification (MLPA) test in a TSC patient revealed a heterozygous deletion spanning exons 1–16 of TSC2. We hence prepared mock samples by mixing genomic DNA from this patient and the patient’s mother at a 1:9 ratio. The exact patient DNA fraction (10.8%) could be measured by multiplex genotyping dPCR^17^. We then designed a dPCR assay targeting the deleted region in *TSC2* and another assay targeting Chr 1 as reference (Supplementary Table 9). We conducted five independent measurements on the mock samples as well as negative controls, and all of the results agreed with the corresponding genotypes using a likelihood ratio classifier (Fig. 4f-g, Methods and Supplementary Table 10). These findings demonstrate the readiness of CLEAR-dPCR for further clinical assay development.

## Discussion

Based on comparisons with existing dPCR methods, CLEAR-dPCR has three major advantages. First, CLEAR-dPCR is a lossless approach. Microfluidic droplet dPCR methods typically lose >30% of the sample during the compartmentalization and detection steps. In CLEAR-dPCR, centrifugal force guarantees that the aqueous phase is completely transformed into droplets through MiCA centrifugation (Supplementary Fig. 1). Optical clearing and *in situ* 3D imaging through light-sheet illumination ensures thorough interrogation of the entire volume of the emulsion. Using CLEAR-dPCR, true absolute counting is intrinsically achievable with no need for extrapolation to compensate for material loss, and lossless interrogation can further reduce measurement uncertainty (Supplementary Fig. 7). These advantages are most crucial when the nucleic acids being interrogated are precious and low in abundance. Second, CLEAR-dPCR is contamination-free. For most clinical applications that repeatedly amplify specific regions, exposure of the post-amplification products to the laboratory environment poses a great risk for contamination of subsequent operations. In CLEAR-dPCR, once the sample is added into the PCR tube on top of the MiCA, the lid can be closed for good. All subsequent steps including compartmentalization, thermal cycling, and signal detection are performed with the lid closed and the emulsion fully sealed by oil. Third, CLEAR-dPCR is robust and easy to operate. The use of conventional equipment, such as a centrifuge and standard PCR machine, renders CLEAR-dPCR a simple experiment. Optical clearing can be achieved with a pre-mixed reaction buffer and the total hands-on time is less than 1 min per sample, with the total time being less than an hour for dPCR assays. The samples are well sealed, allowing easy operation of the reading instrument, omitting and fluidic manipulation. Moreover, the CLEAR-dPCR emulsion remains intact for days and allows repeated scanning (Supplementary Fig. 12).

In summary, CLEAR-dPCR integrates MiCA-facilitated emulsion generation, betaine-mediated emulsion clearing, and light-sheet illumination imaging to greatly simplify the dPCR experimental process. CLEAR-dPCR achieves various improvements over existing dPCR, including complete compartmentalization from bulk solution to droplet; comparable dynamic range with qPCR; elimination of false-positives through fully sealed operation; and fast digital counting at single-copy resolution level without sample loss. We envision that this simple and inexpensive method will lower the technical and economic barriers of dPCR and have broad application in biology and medicine.

## Methods

### Refractive index matching via addition of certain reagents

Transferring post-amplification PCR product is time- and labor-consuming. Furthermore, it rises the risk of contaminating subsequent assays. However, due to the light scattering at the interface of droplet and carrier fluid, it is very difficult to extract fluorescence signals directly from the emulsion containing massive densely-packed droplets. In order to read the intact PCR emulsion at a high throughput, we develop a refractive index matching strategy to produce optically clear PCR emulsion for parallel light-sheet excitation. Considering usually a big refractive index (RI) difference between water and oil, on one hand, we seek suitable oil with relatively low RI close to the water-phase reagent. Among a few available candidates, silicone is selected for its lower RI value (from 1.37 to 1.41), thus much applicable in this case compared to fatty esters (1.43-1.46), aromatic hydrocarbon (higher than 1.50) and liquid fatty hydrocarbons (higher than 1.40). In our demonstration, we choose silicone DMS-T01.5 (Gelest, Pennsylvania) with low refractive index (1.3880) and viscosity (1.5 cSt), as the base oil. Silicon-based surfactant ES5612 (Evonik, Germany) (5% w/w) is added to the base silicone to produce the final emulsion oil, with refractive index being 1.3903.

On the other hand, the refractive index of original PCR reaction mix (∼1.34) is only slightly higher than that of pure water (1.33). Thus, we need to tune the ingredients of the PCR reaction mix, and hence to increase its refractive index to a value close to 1.39. In practice, we add betaine into the PCR reaction mix to reduce this RI difference. Compared to other commonly used PCR additives, such as glycerol, dimethylsulfoxide (DMSO), tetramethylammonium chloride (TMAC), formamide, and bovine serum albumin (BSA) that usually work at low concentration, betaine has a higher applied concentration up to several molar, which can induce a large tunable range of RI. Furthermore, as a zwitterion quaternary amino acid, betaine is neutral in electrical charge and thereby will not destabilize the emulsion. We verified that through the addition of betaine with 3 to 3.2 M concentration (depending on the concentration of other solutes), a highly clear PCR emulsion with well-matched RI can be successfully prepared for high-speed light-sheet readout (Fig. 2). We also note that bulk PCR reactions with so high concentration of betaine in regular microcentrifuge tubes barely take place, but if the reaction mix is partitioned in a digital format, the amplification works (see Supplementary Methods). We hypothesize that as the amplicons are more concentrated in the picoliter droplets, they can trigger the chain reaction of subsequent circles more effectively.

### Emulsification

In our previous work^15^, we used micro-array channels to generate monodisperse droplet for digital PCR. Here we increase the number of the through-hole channels from 7 to 37, to compensate for the loss of droplet generation rate due to higher viscosity. The reaction mixes are added into the well on the MiCA and centrifuged for 4 min at 15,000 relative centrifuge force (rcf). Around 443 thousand droplets are generated in 200 μL regular PCR microcentrifuge tubes with an average diameter of 41 μm. The transparency of the generated emulsion varies with the concentration of betaine addition (Fig. 2).

### In situ 3D imaging of emulsion using LSFM

Light-sheet fluorescence microscopy (LSFM) has recently emerged as a technique of choice that can image large samples with relatively low phototoxicity and at high speed. Here we build a small-format, macro-view light-sheet fluorescence microscope to noninvasively read the fluorescent signals of massive droplets that stack inside the sealed PCR tube. In our experiment, 488 and 532 nm light-sheets with thickness around 22 μm (1/e^2^ value) are generated to optically section the CLEAR emulsion with FAM and HEX/VIC fluorescence, respectively. The 200-μl centrifuge tube (Axygen, U.S.A.) containing dPCR emulsion inside is submerged into a rectangular glass cuvette filled with silicone oil (mixed by two kinds of raw silicone oil, one with high viscosity and RI, the other low, final RI = 1.390), to minimize the light deflection by the conical plastic tube. Then the droplets stacked in the tube is quickly scanned across the stationary laser light-sheet by a motorized actuator (KSTM25E/M, Thorlabs, U.S.A.). A 2× wide-field detection path consecutively collects the fluorescence signals from the light-sheet illuminated planes of the emulsion. An sCMOS camera (ORCA®-Flash 4.0 V2, Hamamatsu, Japan) sequentially records the fluorescence images with a fine step size of 6.5 μm. For each fluorescence channel, 560 plane images (z depth 3.6 mm) of 550×700 pixels are captured within ∼6 seconds. Then by simply stacking the 560 planes together, the fluorescence distribution of entire clear emulsion can be reconstructed at ∼6.5 μm lateral and ∼22 μm axial resolution that are sufficient to resolve single droplets in three dimensions (Supplementary Fig. 5). Finally, the two-color visualization of the emulsion can be readily created by merging the two-channel results together.

### Image-based droplets counting

We develop a computation procedure to process the raw LSFM image stacks, counting the positive droplets at high accuracy. We first correct the vertical background variance induced by Gaussian laser illumination in the vertical direction and remove the stripe noise caused by the laser diffraction in the Fourier space (see Supplementary Methods). Then we search for intensity local maxima in the 3D image matrices and put down their coordinates in 3 dimensions (Fig. 3c and Supplementary Fig. 6). The intensity pattern around the local maxima are compared with a positive droplet intensity template and assessed by resemblance to filter out bright artefacts. Finally, the positive droplets are located after setting an appropriate threshold between the maxima values and their difference with local backgrounds.

### Duplex CLEAR dPCR for prenatal test of TSC

Peripheral blood DNA samples were harvested from a child patient affected with TSC and a healthy volunteer, whose weight/volume concentrations were first quantitated by fluorometer (Qubit, ThermoFisher, US) as a rough reference. We then set the threshold of presumption of an affected fetus to be 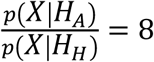, whereas that of a healthy fetus 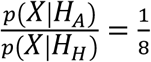. Values between the two thresholds were thought to be invalid. Pure healthy and affected samples were first test by CLEAR dPCR for validation of the assay as well as to further quantify the DNA concentration. Healthy DNA sample doped with the affect was used as mock maternal cell free DNA of a mother carrying a affected fetus (fetal fraction = 10.8%). Each independent assay consisted six tubes for CLEAR-dPCR so that the DNA counts were enough to give a solid result. ^17^ We carried 5 parallel assays and the results came as expected, all being positive. (Fig. 4f,g and Supplementary Table 9,10).

## Acknowledgments

We thank Dr. Wenxiong Zhou for fruitful discussion. This work was supported by Ministry of Science and Technology of China (2018YFA0108100 to Y.H. and 2017YFA0700500 to P.F.), National Natural Science Foundation of China (21525521 to Y.H., 21874052 to P.F., and 21675098 to J.W.), the Innovation Fund of WNLO, and Beijing Advance Innovation Center for Genomics.

## Author Contributions

Y.H., P.F., J.W. and Z.C. conceived and oversaw the project. P.L., F.Z., and Y.S. developed the clear emulsion protocol. M.J. and J.N. developed optical setup. P.L. and M.D. designed the molecular assays. P.L. and M.J. conducted the dPCR experiments and processed the images. P.L., M.J., Z.C., J.W., P.F., and Y.H. analyzed the data and wrote the paper.

## References

1. Wong, Y. K. et al. Applications of digital PCR in precision medicine. Expert Review of Precision Medicine and Drug Development 2, 177–186 (2017).

2. Olmedillas-López, S., García-Arranz, M. & García-Olmo, D. Current and Emerging Applications of Droplet Digital PCR in Oncology. Molecular Diagnosis & Therapy 21, 493–510 (2017).

3. Robin, J. D., Ludlow, A. T., LaRanger, R., Wright, W. E. & Shay, J. W. Comparison of DNA Quantification Methods for Next Generation Sequencing. Sci Rep 1–10 (2016). doi:10.1038/srep24067

4. Vogelstein, B. & Kinzler, K. W. Digital PCR. Proc Natl Acad Sci USA 96, 9263–9241 (1999).

5. Sykes, P. J. et al. Quantitation of Targets for PCR by Use of Limiting Dilution. Biotechniques 13, 444–449 (1992).

6. Ottesen, E. A., Hong, J. W., Quake, S. R. & Leadbetter, J. R. Microfluidic digital PCR enables multigene analysis of individual environmental bacteria. Science (New York, N.Y.) 314, 1464–1467 (2006).

7. Heyries, K. A. et al. Megapixel digital PCR. Nat Meth 8, 649–651 (2011).

8. Shen, F., Du, W., Kreutz, J. E., Fok, A. & Ismagilov, R. F. Digital PCR on a SlipChip. Lab Chip 10, 2666–2672 (2010).

9. Hatch, A. C. et al. 1-Million droplet array with wide-field fluorescence imaging for digital PCR. Lab Chip 11, 3838–3845 (2011).

10. Lo, Y. M. et al. Digital PCR for the molecular detection of fetal chromosomal aneuploidy. Proceedings of the National Academy of Sciences 104, 13116–13121 (2007).

11. Dangla, R., Kayi, S. C. & Baroud, C. N. Droplet microfluidics driven by gradients of confinement. Proc Natl Acad Sci U S A 110, 853–858 (2013).

12. Kreutz, J. E. et al. Theoretical design and analysis of multivolume digital assays with wide dynamic range validated experimentally with microfluidic digital PCR. Anal. Chem. 83, 8158–8168 (2011).

13. Peiyu LiaoYanyi Huang. Digital PCR: Endless Frontier of ‘Divide and Conquer’. Micromachines 8, 231–7 (2017).

14. Baker, M. Digital PCR hits its stride. Nat. Methods 9, 541–544 (2012).

15. Chen, Z. et al. Centrifugal micro-channel array droplet generation for highly parallel digital PCR. Lab Chip 17, 235–240 (2017).

16. Henke, W., Herdel, K., Jung, K., Schnorr, D. & Loening, S. A. Betaine improves the PCR amplification of GC-rich DNA sequences. 1–2 (1997).

17. Camunas-Soler, J. et al. Noninvasive Prenatal Diagnosis of Single-Gene Disorders by Use of Droplet Digital PCR. Clinical Chemistry 64, 336–345 (2018).

18. Kuliev, A. & Rechitsky, S. Preimplantation genetic testing: current challenges and future prospects. Expert Review of Molecular Diagnostics 17, 1071–1088 (2017).

19. Willis, A. S., Veyver, I. & Eng, C. M. Multiplex ligation-dependent probe amplification (MLPA) and prenatal diagnosis. Prenat Diagn 32, 315–320 (2012).

20. Jones, A. C. et al. Comprehensive Mutation Analysis of TSC1 and TSC2- and Phentypic Correlations in 150 Families with Tuberous Sclerosis. American Journal of Human Genetics 64, 1305–1315 (1999).

21. Martin, K. R. et al. The genomic landscape of tuberous sclerosis complex. Nature Communications 8, 1–13 (2017).

