## Supplementary Information for "Lossless and Contamination-Free Digital PCR"

**Table of Contents**

I. Supplementary Figures (Fig. 1-12)
II. Supplementary Tables (Table 1-10)
III. Supplementary Methods
1. Betaine changes water solution's refractive index
2. CLEAR dPCR assay
    2.1 DNA sequences
    2.2 Protocol
3. MiCA emulsification free of DNA adsorption
4. The optical setup of readout system
    4.1 The light path
    4.2 Components of the readout system
    4.3 Characterization of the setup
5. 3D image-based droplet counting
6. Results of CLEAR PCR versus those of QX200 ddPCR
7. Light-sheet illumination at low photobleaching rate supports repetitive CLEAR dPCR imaging
8. PCR not interfered by high concentration of betaine in CLEAR dPCR emulsion
9. Lossless dPCR reduces quantitative uncertainty

### I. Supplementary Figures

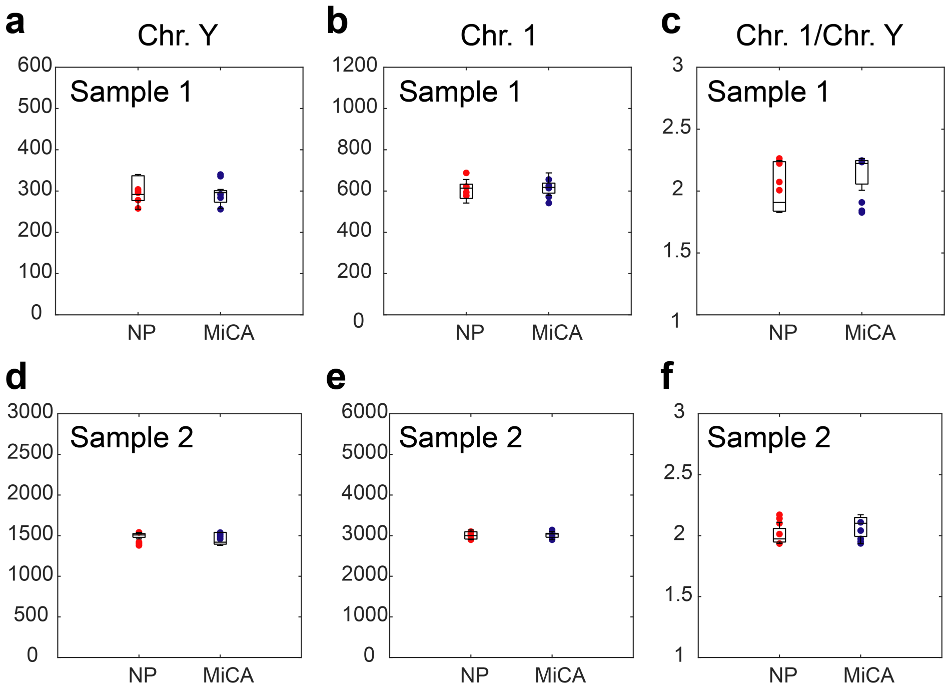

**Supplementary Fig. 1 | Compartmentation through MiCA does not cause sample loss**. Copy number quantification was done using Bio-Rad QX200 digital PCR machine as an orthogonal validation. Genomic DNA aliquots, with (blue dots) and without (red dots, 'NP' for 'not processed') MiCA centrifugation treatment, are subject to quantification. The copy number-ratio between chromosome 1 and chromosome Y is 2, indicating the male-origin of the genomic DNA. We tested gDNA samples in two different concentrations 3.1ng (**a**-**c**) and 15.7 ng (**d**-**f**), respectively.

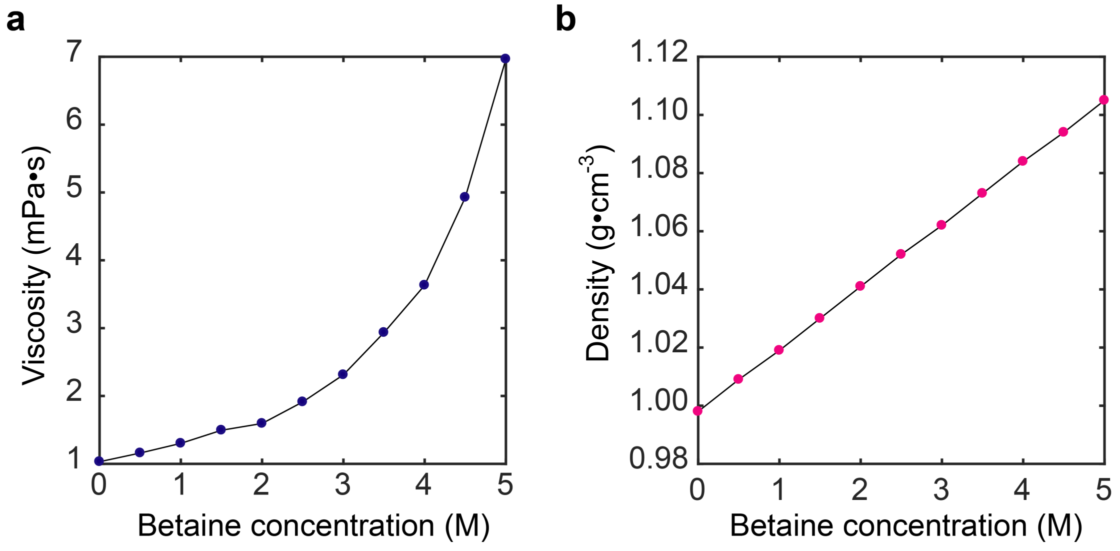

**Supplementary Fig. 2 | Two major physical properties**. Viscosity (**a**) and density (**b**), are dependent upon the concentration of betaine in water. Both viscosity and density are elevated with the addition of betaine.

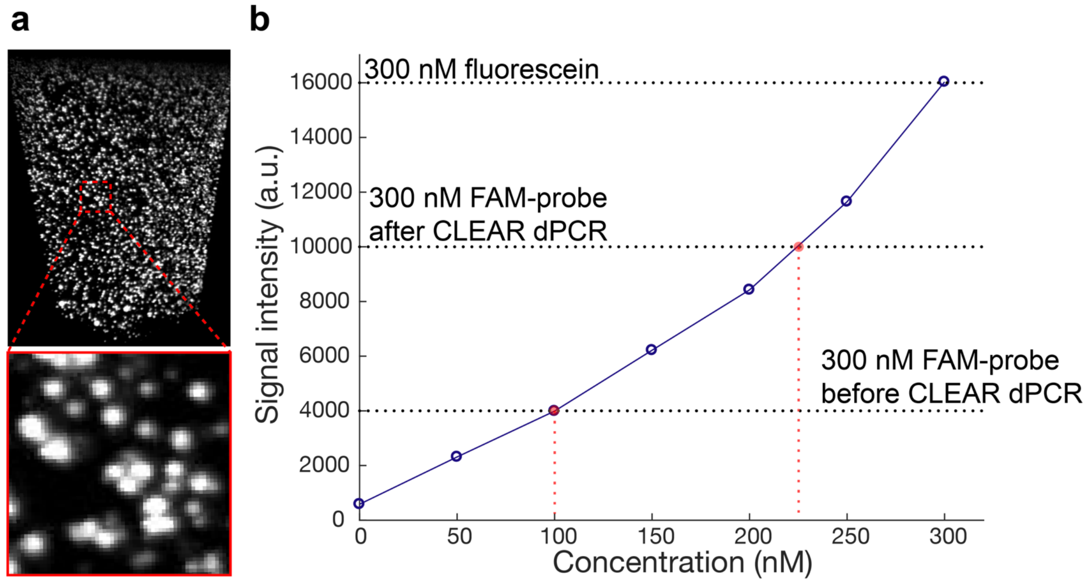

**Supplementary Fig. 3 | Assessment of the hydrolyzed ratio of hydrolysis probes used in CLEAR-dPCR experiments that containing high concentration betaine**. **a**, Light-sheet excited fluorescence image of mixture of 3.14-M betaine containing droplets with and without fluorescein, to mimic the CLEAR-dPCR emulsion. **b**, With different fluorescein concentration in the droplets, a standard curve for intensity-concentration calibration is obtained. The droplet fluorescence intensities of hydrolysis-probe before and after CLEAR dPCR have also been shown.

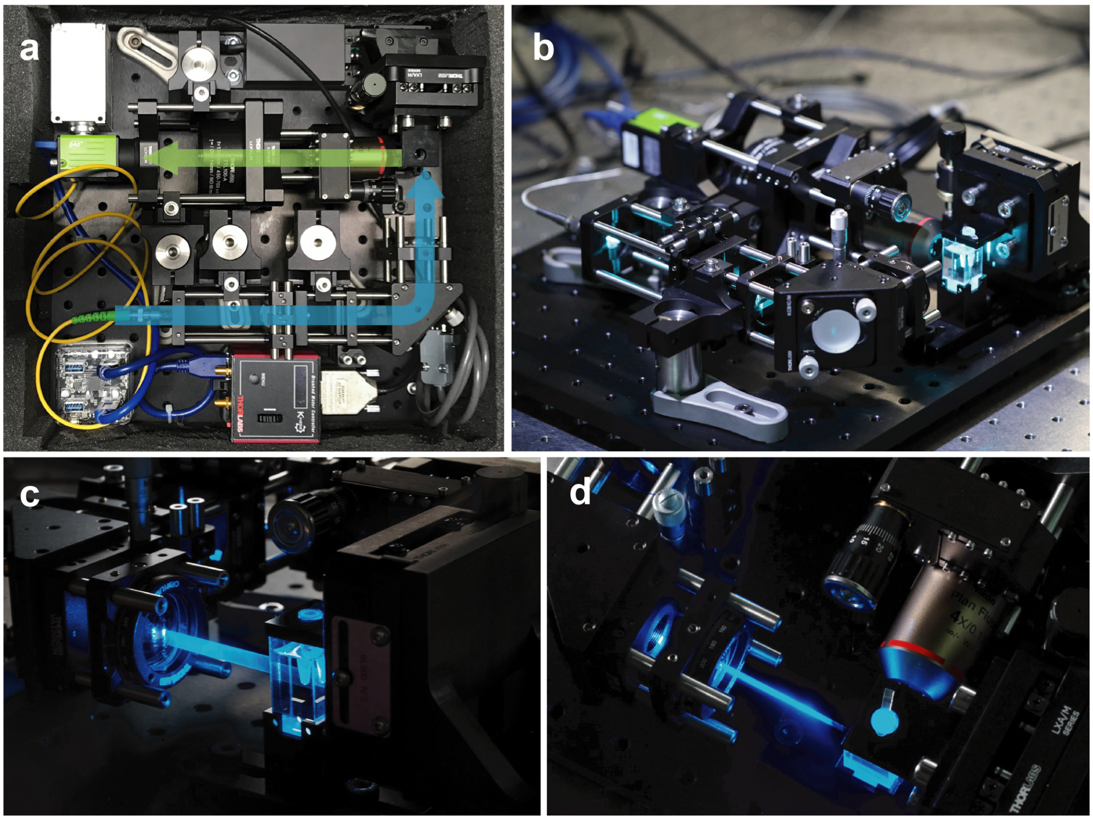

**Supplementary Fig. 4 | The lab-built light-sheet dPCR 3D reader, constructed from common opto-mechanical parts**. **a**, Top view of the setup, with excitation light path (blue) and fluorescence collection light path (green) labeled. **b**, The whole setup, not including the laser head and laser power unit, is compact, sitting on a 30 x 30 cm breadboard. **c-d**, Optical sectioning of the PCR tube by a light sheet.

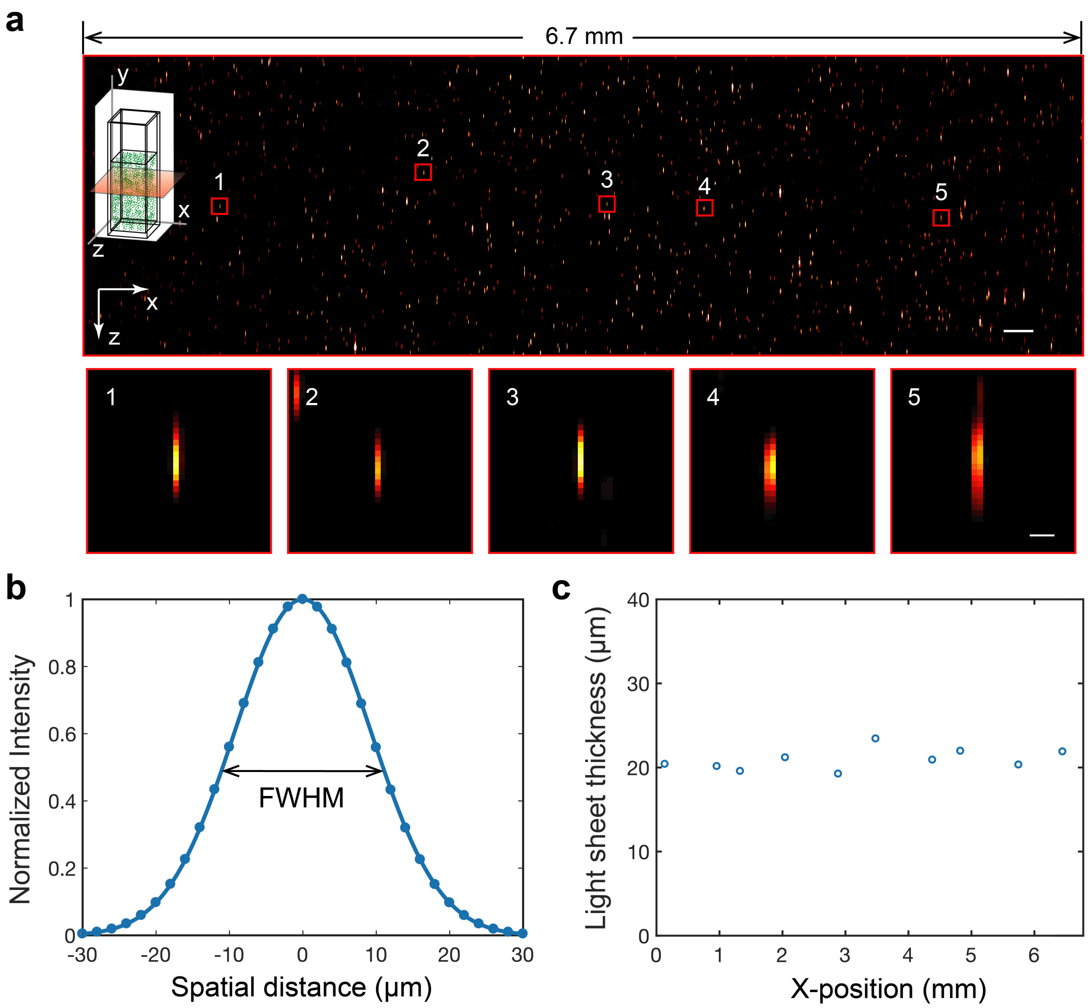

**Supplementary Fig. 5 | Characterization of the light-sheet**. **a**, The reconstructed x-z plane of the sub-resolution fluorescent beads (diameter 1 μm). Five beads located in different position of the image are shown in vignette high-resolution views. Scale bars: big field image, 200 μm; vignettes, 10 μm. **b**, The intensity profile of the resolved bead along z direction, with FWHM about 22 μm. **c**, Light sheet thickness, measured by the beads FWHM, along the x-axis (span ~6.7 mm).

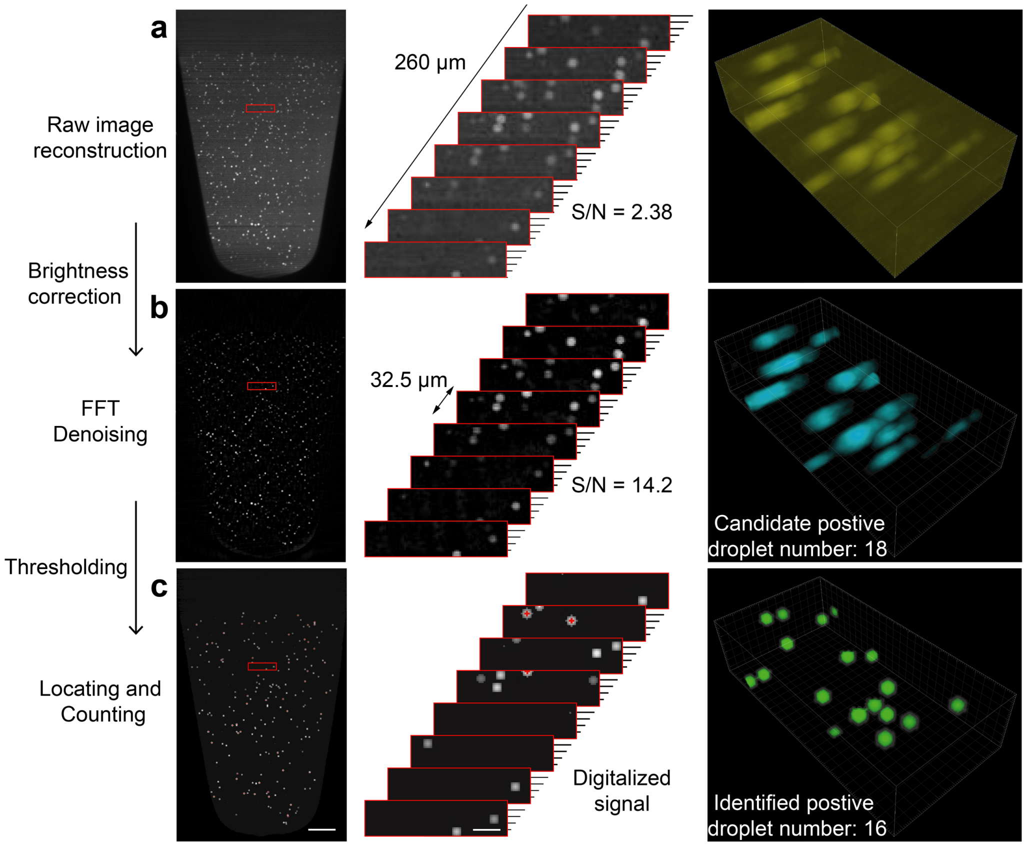

**Supplementary Fig. 6 | Image processing and 3D counting in CLEAR dPCR**. **a**, The raw images taken from sCOMS camera were reconstructed into 3D stacks. **b**, After intensity profile correction and denoising, the signal-to-noise ratio of images were improved. **c**, Through further thresholding, the positive droplets were identified. (Scale bars: big field image, 500 µm; vignettes, 100 µm)

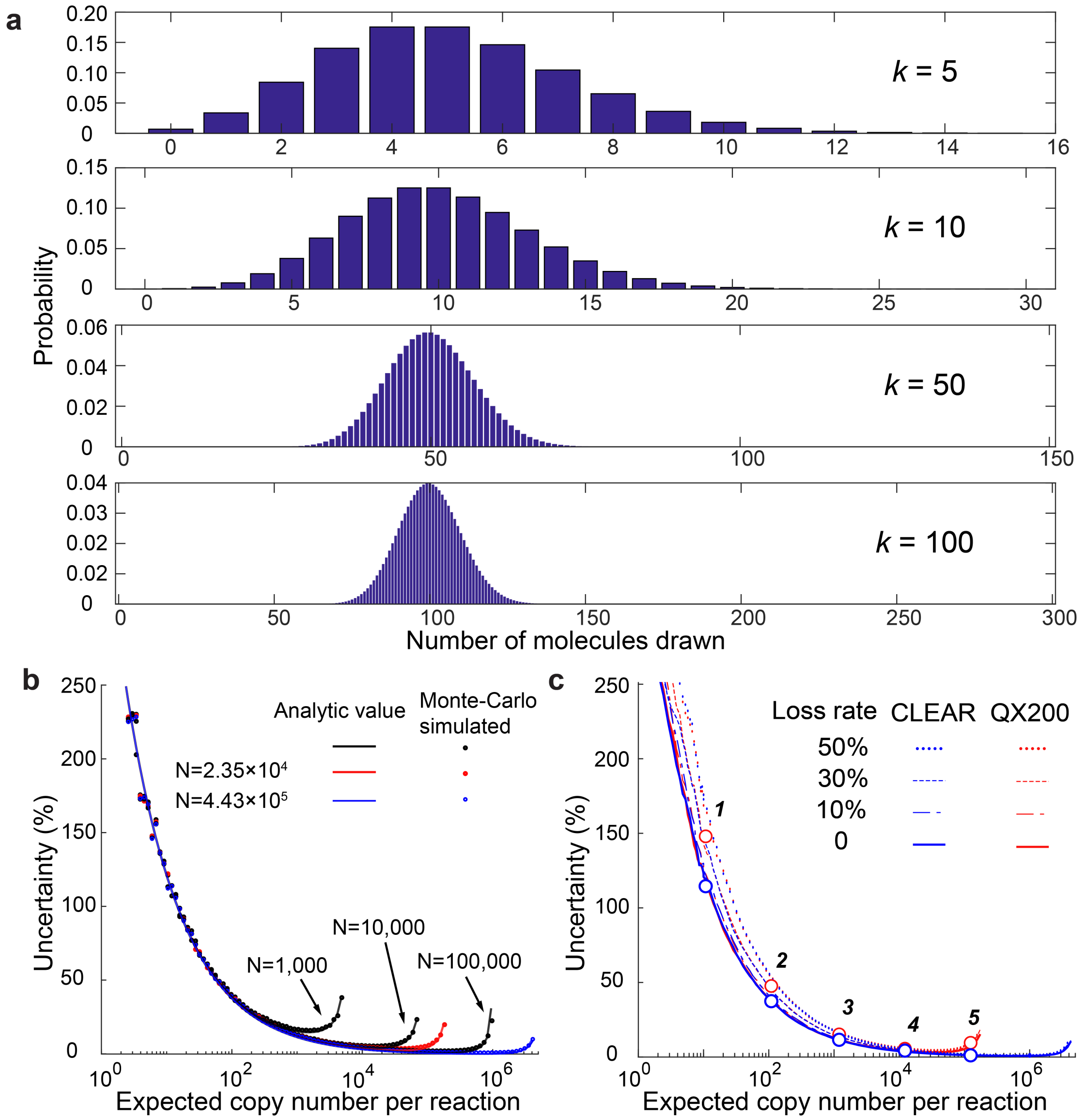

**Supplementary Fig. 7 | Uncertainty in digital PCR**. **a**, Subsampling induced uncertainty, reflected as a probability distribution, with different expected number of molecules of each reaction according to Poisson distribution's probability mass function. **b**, Overall uncertainty (subsampling and partitioning combined) in dPCR with different partition numbers (*N*). Both analytical analysis and Monte Carlo simulation (10,000 repeats) have been carried and their results agree well with each other. Without droplet loss, increasing the compartmentation number will extend the detection range with low uncertainty. We also plot two specific curves for Bio-Rad QX200 ddPCR system (red) and for CLEAR dPCR (blue). With 20-fold more compartments, CLEAR dPCR allows quantifying high concentration samples with high accuracy. **c**, The Monte Carlo simulation on the digital

PCR uncertainty at different rate of partition loss. In an actual experimental practice, any loss of counting partition will lead to a high uncertainty.

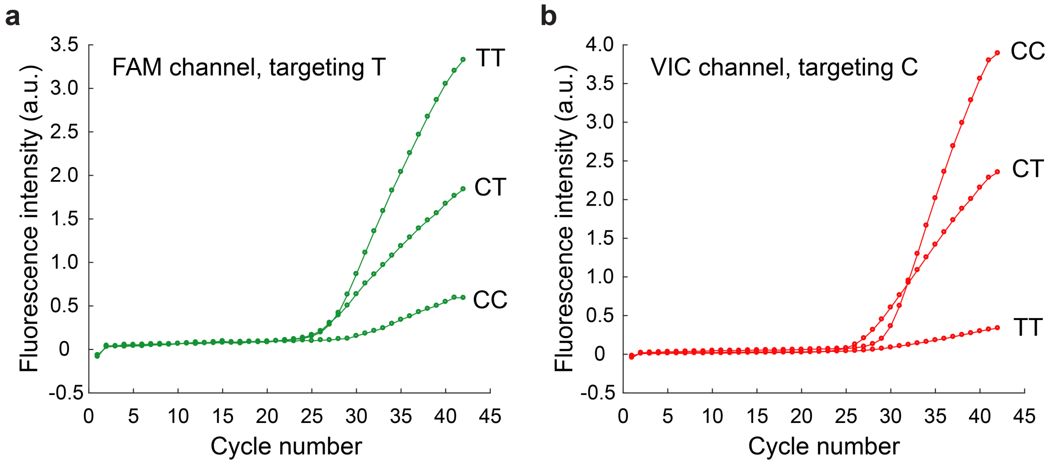

**Supplementary Fig. 8 | Real-time quantitative PCR validation of self-design hydrolysis probes for SNP genotyping**. Genomic DNA samples of genotypes CC, CT, and TT can be well differentiated.

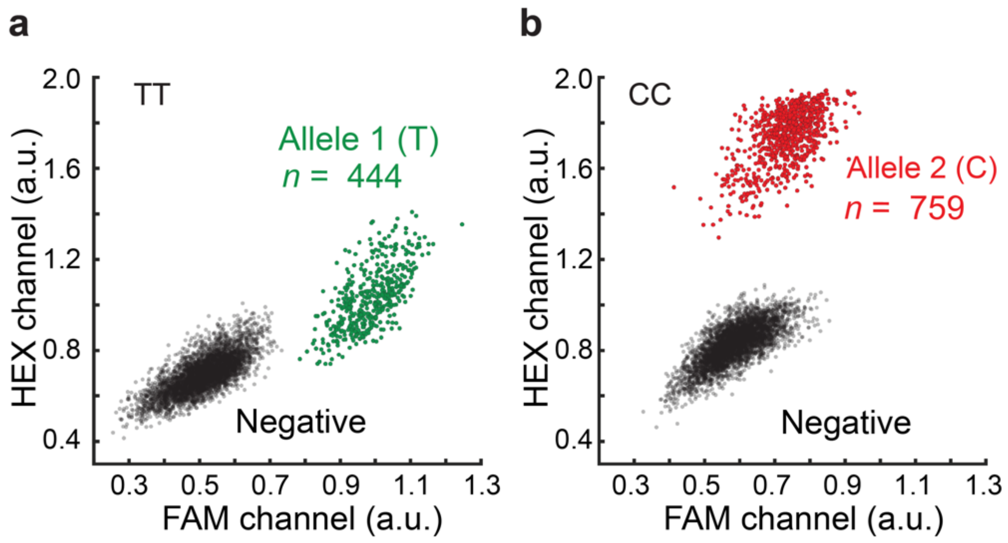

**Supplementary Fig. 9 | SNP detection**. Scatterplot results of samples that are homozygous at SNP rs10092491.

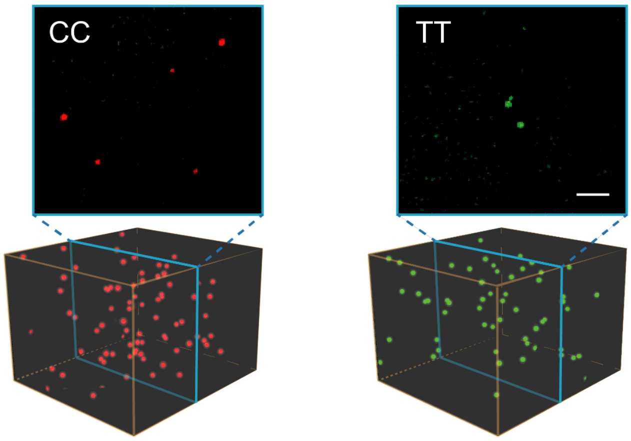

**Supplementary Fig. 10 | 3-D signal reconstructions of the 2 homozygous samples (CC and TT).** As shown, the signal crosstalk between channels are negligible. Scale bar: 200 μm.

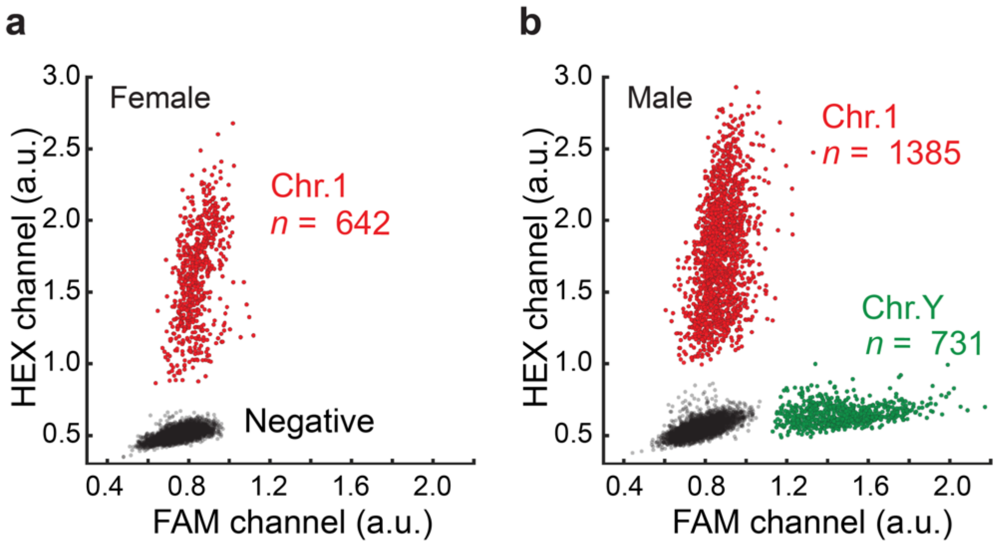

**Supplementary Fig. 11 | Copy number variation detection by CLEAR-dPCR**. Copy numbers of chromosomes 1 (red) and Y (green) were detected in a female and male human genomic DNA samples.

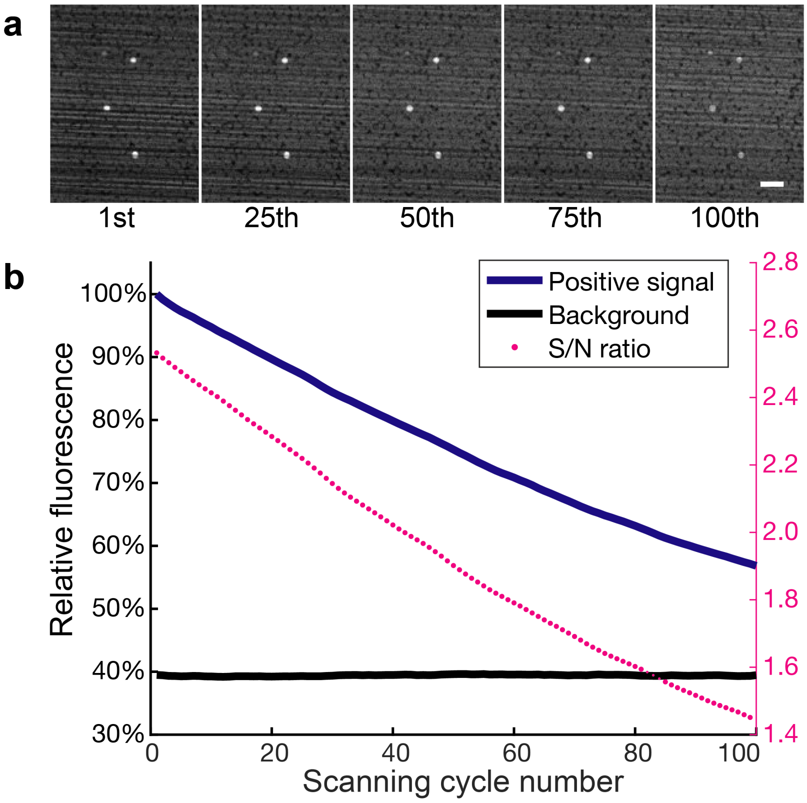

**Supplementary Fig. 12** | Photostability of the fluorescent droplets. **a**, The raw images of CLEAR dPCR samples repetitively scanned at $1^{st}$, $25^{th}$, $50^{th}$, $75^{th}$, and $100^{th}$ cycles. Scale bar: 200 μm. **b**, Normalized fluorescence intensity plot of positive signal, background, and signal-to-noise (S/N) ratio versus the number of scanning cycles.

#### II. Supplementary Tables

**Supplementary Table 1 | DNA sequences used in absolute quantification assay.**

| Oligo-DNAs | Sequence |
|---|---|
| prfA gene<br>280-bp fragment<br>(template) | **CCGCAAATAGAGCCAAGCTT**CCCGTTAATCGAAAAATCATTAAAT<br>TTAGCTAGACTGTATGAAACTTGTTTTTGTAGGGTTTGGAAAACA<br>TAGAAAAAGTGCGTAAGATTCTTGCTCAGTAGTTCTTTTAGTTCG<br>TTTATTTTGATAACGTATGCGGTA**GCCTGTTCGCTAATGACTTCT**<br>**AAAT**TA**TAATAGCCAACCGATGTTT**CTGTATCAAT**AAAGCCAGAC**<br>**ATTATAACGAAAGCAC**CTTTGTAGTATTGTAAATT**CATGATGGTC**<br>**CCGTTCTCAC** |
| Forward primer for<br>template<br>Production | 5′-**CCGCAAATAGAGCCAAGCTT**-3′ |
| Reverse primer for<br>template production | 5′-**GTGAGAACGGGACCATCATG**-3′ |
| Forward primer for<br>dPCR | 5′-**GCCTGTTCGCTAATGACTTCTAAAT**-3′ |
| Reverse primer for<br>dPCR | 5′-**GTGCTTTCGTTATAATGTCTGGCTTT**-3′ |
| Probe | 5′-FAM-**TAATAGCCAACCGATGTTT**-MGB-3′ |

**Supplementary Table 2 | Absolute quantification by CLEAR dPCR**.

| Samples | | Counted copy number in 16 μl | | | | | Avg. count | DNA copy No. CV |
|---|---|---|---|---|---|---|---|---|
| | | A | B | C | D | E | | |
| Neg (-) Ctrl. | Pos (+) droplet | 0 | 0 | 0 | 0 | 0 | 0 | 0.0% |
| | DNA Copies* | 0 | 0 | 0 | 0 | 0 | 0 | |
| Conc. 1 | Pos (+) droplet | 10 | 8 | 11 | 13 | 17 | 11.8 | 25.9% |
| | DNA Copies | 10.0 | 8.0 | 11.0 | 13.0 | 17.0 | 11.8 | |
| Conc. 2 | Pos (+) droplet | 125 | 115 | 134 | 133 | 128 | 127.0 | 5.4% |
| | DNA Copies | 125.0 | 115.0 | 134.0 | 133.0 | 128.0 | 127.0 | |
| Conc. 3 | Pos (+) droplet | 1304 | 1175 | 1241 | 1214 | 1206 | 1228.0 | 3.5% |
| | DNA Copies | 1305.9 | 1176.6 | 1242.7 | 1215.7 | 1207.6 | 1229.7 | |
| Conc. 4 | Pos (+) droplet | 12100 | 12295 | 12216 | 13688 | 13044 | 12668.6 | 4.9% |
| | DNA Copies | 12268.3 | 12468.8 | 12387.6 | 13903.9 | 13239.9 | 12853.7 | |
| Conc. 5 | Pos (+) droplet | 108462 | 110005 | 106501 | 109687 | 110006 | 108932.2 | 1.4% |
| | DNA Copies | 124402.9 | 126450.9 | 121813.7 | 126028.1 | 126452.3 | 125029.6 | |

*DNA copy number = $-N \times \ln(1 - N_p/N)$, where $N$ is the total droplet number and $N_p$ are positive count, and in this assay we assign $N = 4.43 \times 10^5$.

#### Supplementary Table 3 | Digital PCR assay on QX200.

| Samples | | Estimated copy number in 20 µl | | | | | Avg. count | Conc. CV |
|---|---|---|---|---|---|---|---|---|
| | | A | B | C | D | E | | |
| Neg (-) Ctrl. | Pos (+) count | 0 | 0 | 0 | 0 | 0 | 0.0 | 0.0% |
| | Neg (-) count | 12278 | 12041 | 12308 | 11185 | 11007 | | |
| | DNA Copies | 0 | 0 | 0 | 0 | 0 | | |
| Conc. 1 | Pos (+) count | 6 | 3 | 5 | 4 | 4 | 6.12 | 23.6% |
| | Neg (-) count | 16879 | 16958 | 16594 | 17233 | 17041 | | |
| | DNA Copies | 8.4 | 4.2 | 7 | 5.4 | 5.6 | | |
| Conc. 2 | Pos (+) count | 92 | 79 | 97 | 93 | 77 | 124.80 | 6.3% |
| | Neg (-) count | 16594 | 16529 | 17731 | 16219 | 15056 | | |
| | DNA Copies | 130 | 112 | 128 | 134 | 120 | | |
| Conc. 3 | Pos (+) count | 884 | 860 | 1026 | 834 | 821 | 1292.00 | 5.3% |
| | Neg (-) count | 15562 | 15307 | 16588 | 15871 | 14978 | | |
| | DNA Copies | 1300 | 1286 | 1412 | 1206 | 1256 | | |
| Conc. 4 | Pos (+) count | 6319 | 6443 | 4407 | 6530 | 6038 | 12508.00 | 2.3% |
| | Neg (-) count | 8831 | 9679 | 6092 | 9134 | 8741 | | |
| | DNA Copies | 12700 | 12000 | 12800 | 12680 | 12360 | | |
| Conc. 5 | Pos (+) count | 15492 | 15255 | 13781 | 14519 | 13158 | 142000.00 | 4.2% |
| | Neg (-) count | 32 | 503 | 28 | 31 | 49 | | |
| | DNA Copies | 145600 | 81000 | 146000 | 144800 | 131600 | | |

**Supplementary Table 4 | Real time qPCR for absolute quantification.**

| Samples | | Estimated copy number in 20 µl | | | | | Avg. | Conc. |
|---|---|---|---|---|---|---|---|---|
| | | A | B | C | D | E | Count | CV |
| Neg (-) Ctrl. | Ct No. | 39.76 | 39.35 | 39.76 | 38.41 | 40.39 | 39.53 | -/- |
| | Copies/20 µl* | -/- | -/- | -/- | -/- | -/- | -/- | |
| Conc. 1 | Ct No. | 36.75 | 37.46 | 36.88 | 37.19 | 37.04 | 37.07 | 2.3% |
| | Copies/20 µl | 16.4 | 10.4 | 15.1 | 12.4 | 13.6 | 13.6 | |
| Conc. 2 | Ct No. | 34.09 | 33.95 | 33.79 | 33.70 | 34.01 | 33.90 | 2.4% |
| | Copies/20 µl | 89.5 | 98.4 | 108.9 | 115.1 | 94.6 | 101 | |
| Conc. 3 | Ct No. | 29.84 | 29.84 | 29.67 | 29.72 | 29.68 | 29.75 | 5.4% |
| | Copies/20 µl | $1.36\times10^3$ | $1.36\times10^3$ | $1.51\times10^3$ | $1.46\times10^3$ | $1.51\times10^3$ | $1.44\times10^3$ | |
| Conc. 4 | Ct No. | 26.29 | 26.22 | 26.20 | 26.26 | 26.21 | 26.24 | 10.4% |
| | Copies/20 µl | $1,31\times10^4$ | $1.37\times10^4$ | $1.39\times10^4$ | $1.35\times10^4$ | $1.39\times10^4$ | $1.36\times10^4$ | |
| Conc. 5 | Ct No. | 22.92 | 22.89 | 22.93 | 22.91 | 22.84 | 22.90 | 17.2% |
| | Copies/20 µl | $1.14\times10^5$ | $1.16\times10^5$ | $1.13\times10^5$ | $1.14\times10^5$ | $1.20\times10^5$ | $1.15\times10^5$ | |

*To estimate the DNA count number in each well of 20 µl reaction mix, a standard curve is obtained by linear regression using the five positive sets of assays: DNA copy number = $10^{\left(\frac{41.12-Ct}{3.601}\right)}$.

**Supplementary Table 5 | DNA sequences in duplex assay to differentiate SNP.**

| | | |
|---|---|---|
| SNP of interest | TCTGTGATAGAGTGGCATTAGAAATTCCAGATAGAGCTAAAACTGAAGYTTTCCTTATAGAGATTTATCCTAGTTAGTTTGCGGGG | SNP ID: rs10092491 Location: Chr8:28553555 |
| Fwd. primer | 5′-TCTGTGATAGAGTGGCATTAGAAATTC-3′ | |
| Rev. primer | 5′-CCCCGCAAACTAACTAGGATAAATC-3′ | |
| Probe 1 | FAM-5′-CTAAAACTGAAGCTTTC-3′-MGB | 488-nm channel |
| Probe 2 | VIC-5′-AACTGAAGTTTTCCTTATAG-3′-MGB | 532-nm channel |

#### Supplementary Table 6 | SNP genotyping

| CLEAR dPCR | | | | | |
|---|---|---|---|---|---|
| Sample | Positive Counts | | | Ch1/Ch2 | Result |
| | 488 nm / C | 532 nm / T | Both Channels | | |
| PH01 | 611 | 646 | 4 | 97.5% | CT |
| PC02 | 497 | 0 | 0 | n/a | CC |
| P112 | 357 | 0 | 0 | n/a | CC |
| PX39 | 243 | 224 | 15 | 0.0% | CT |
| PX61 | 0 | 63 | 0 | 0.0% | TT |
| PX76 | 271 | 280 | 13 | 108.9% | CT |
| PX79 | 0 | 246 | 0 | 0.0% | TT |
| PH10 | 818 | 884 | 7 | 92.5% | CT |

| QX200 ddPCR | | | | | | |
|---|---|---|---|---|---|---|
| Sample | Positive Counts | | | | Ch1/Ch2 | Result |
| | Chn. 1 / C | Chn. 2 / T. | Both Chn. | Negative | | |
| PH01 | 100 | 93 | 1 | 7605 | 107.5% | CT |
| PC02 | 81 | 0 | 0 | 9073 | n/a | CC |
| P112 | 382 | 0 | 0 | 7858 | n/a | CC |
| PX39 | 93 | 104 | 1 | 11329 | 1.0% | CT |
| PX61 | 0 | 141 | 0 | 13704 | 0.0% | TT |
| PX76 | 230 | 213 | 4 | 13388 | 108.0% | CT |
| PX79 | 0 | 123 | 0 | 9663 | 0 | TT |
| PH10 | 868 | 874 | 64 | 11424 | 99.3% | CT |

**Supplementary Table 7 | DNA sequences in duplex assay for two independent sequences (sex determination).**

| | | |
|---|---|---|
| Target sequence 1 | **AGTTTCGAACTCTGGCACCT**TTCAATTT**TGTCGCACTCTCCTTGTTTTT**GACAATGCAATCATA**TGCTTCTGCTATGTTAAGCGTAT** | Location: chrY:2,787,656-2,787,570 |
| Fwd. primer 1 | 5′-**AGTTTCGAACTCTGGCACCT**-3′ | |
| Rev. primer 1 | 5′-**CGGTTAACATAGCAGAAGCA**-3′ | |
| Probe 1 | FAM-5′-**TGTCGCACTCTCCTTGTTTTT**-3′-IBFQ* | 488-nm channel |
| Target sequence 2 | **GTTCGGCTTTCACCAGTCT**GTG**CGCCCTGCCATGTGGAAGA**TGATGCTCAACATTGATGGT**GAGTGGGGAGAGCTATGGAG** | Location: chr1:36,475,730-36,475,810 |
| Fwd. primer 2 | 5′-**GTTCGGCTTTCACCAGTCT**-3′ | |
| Rev. primer 2 | 5′-**CTCCATAGCTCTCCCCACTC**-3′ | |
| Probe 2 | HEX-5′-**CGCCCTGCC**/ZEN/†**ATGTGGAAGA**-3′-IBFQ* | 532-nm channel |

*IBFQ is Iowa Back fluorescent quencher. It is from Integrated DNA Technologies (IDT).

†ZEN is a second quencher that is added at the middle of probes for lower detection background, also from IDT.

#### Supplementary Table 8 | Sex determination assays.

| CLEAR dPCR | | | | | | |
|---|---|---|---|---|---|---|
| Sample | Positive Counts | | | | Ch1/Ch2 | Result |
| | 488 nm / chrY | 532 nm / chr1 | Both Channels | Negative | | |
| PH01 | 577 | 1202 | 2 | NA | 48.0% | M |
| PC02 | 282 | 521 | 4 | NA | 54.1% | M |
| P112 | 800 | 1445 | 6 | NA | 55.4% | M |
| P113 | 343 | 644 | 2 | NA | 53.2% | M |
| PX57 | 0 | 624 | 0 | NA | 0.0% | F |
| PX61 | 0 | 881 | 0 | NA | 0.0% | F |
| PX76 | 0 | 862 | 0 | NA | 0.0% | F |
| PX79 | 0 | 707 | 0 | NA | 0.0% | F |
| QX200 ddPCR | | | | | | |
| Sample | Positive Counts | | | | Ch1/Ch2 | Result |
| | Chn.1 / chrY | Chn. 2 / chr1 | Both Channels | Negative | | |
| PH01 | 124 | 262 | 2 | 10050 | 47.3% | M |
| PC02 | 324 | 689 | 21 | 10791 | 47.0% | M |
| P112 | 251 | 540 | 12 | 10987 | 46.5% | M |
| P113 | 102 | 203 | 1 | 9314 | 50.2% | M |
| PX57 | 0 | 183 | 0 | 7436 | 0.0% | F |
| PX61 | 0 | 387 | 0 | 12818 | 0.0% | F |
| PX76 | 1 | 325 | 0 | 11220 | 0.4% | F |
| PX79 | 0 | 446 | 0 | 11537 | 0.0% | F |

**Supplementary Table 9 | DNA sequences in duplex assay for two independent sequences (TSC2 assay).**

| | | |
|---|---|---|
| Target sequence 1 | GCTTCCGCATGACTTTGGAGGACCGCATTAGTC<br>GAGTCTGTTAACCACAGCTTTAAGGAGGACTCC | Location:<br>chr16:2,061,147-2,061,212 |
| Fwd. primer 1 | 5′-GCTTCCGCATGACTTTGGA-3′ | |
| Rev. primer 1 | 5′-GGAGTCCTCCTTAAAGCTGTGGTT-3′ | |
| Probe 1 | FAM-5′-ACCGCATTAGTCGAGTCT-3′-MGB | 488-nm channel |
| Target sequence 2 | CCCTGGATGTGGCCATGAGGCACCTGGCATCCA<br>TGAGGTATTGGGTGTAGTTAGTATCTGGG | Location:<br>chr1:35,893,242-35,893,303 |
| Fwd. primer 2 | 5′-CCCTGGATGTGGCCATGA-3′ | |
| Rev. primer 2 | 5′-CCCAGATACTAACTACACCCAATACCT-3′ | |
| Probe 2 | HEX-5′-CACCTGGCATCCAT-3′-MGB | 532-nm channel |

#### Supplementary Table 10 | Prenatal TSC assay

| Fetal fraction determination | | | | |
|---|---|---|---|---|
| Sample | Positive Counts | | 2 × Ch1/Ch2 | w/v concentration |
| | 488 nm / C | 532 nm / T | | |
| H1 | 6685 | 6661 | 2.007 | 1.50 ng/μl |
| A1 | 2713 | 5201 | 1.043 | 1.07 ng/μl |

Based on w/v concentration, we tried to dope 10% affected DNA into the healthy sample, and we mixed 193 μl healthy sample with 30μl the affected. Fetal fraction needs to be corrected due to w/v concentration measurement error.

$$\varepsilon = \frac{C_A V_A}{C_H V_H + C_A V_A} = \frac{Count_{A,488} V_A}{Count_{A,488} V_A + Count_{H,488} V_H} = \frac{5201\times 30}{5201\times 30 + 6661\times 193} \approx 10.8\%$$

| Prenatal TSC assay using mock maternal peripheral DNA | | | | |
|---|---|---|---|---|
| Sample | Positive Counts | | Copy number (2 × Ch1/Ch2) | Positive likelihood* |
| | 488 nm / C | 532 nm / T | | |
| HA1 | 6577 | 6863 | 1.917 | 99.6% |
| HA2 | 5626 | 6045 | 1.861 | 100.0% |
| HA3 | 5374 | 5634 | 1.908 | 99.7% |
| HA4 | 5665 | 5934 | 1.909 | 99.7% |
| HA5 | 5227 | 5593 | 1.869 | 100.0% |

*The positive likelihood $p_{(+)}$ is calculated by the following equation:

$$p_{(+)} = \frac{p\ (X|H_A)}{p\ (X|H_A) + p\ (X|H_H)}$$

Where $p\ (X|H_A)$ is probability coming from an affected fetus and $p\ (X|H_H)$ that from a healthy fetus.

#### III. Supplementary Methods.

##### 1. Adding betaine in water elevates solution's refractive index

With betaine added in the PCR mixes, several physical properties of the solution such as the refractive index (RI), density and viscosity will significantly change. These changes should be carefully considered because they are closely related to the choice of emulsion oil and design of droplet generation protocol. Especially, the RI of solution will be significantly elevated by betaine addition, linearly responsive to the concentration ($\Delta$RI = 0.01665 $mol^{-1}$, with $R$ = 0.9998). Given the betaine's solubility being around 5.2 M, the maximum refractive index of PCR mixes could reach ~1.42, which is within the range of silicones' RI and thus well suited to the generation of optical clear emulsion through RI matching.

It is also noteworthy that as the betaine concentration increases, the viscosity of solution changes dramatically at a rate much higher than the increase of its density. This effect substantially extends the emulsification's duration using MiCA in our previous design (7 holes). To accelerate this process, we increased the number of through-holes to 37, and hence improved the centrifugal speed from 13,000 rcf to 15,000 rcf. Finally, the emulsification could be completed within 4 min.

##### 2. CLEAR dPCR assay

*2.1 DNA Sequences*

In the simplex dPCR for absolute quantification we chose a sequence from Listeria monocytogenes *prfA* gene. The oligonucleotide sequences are listed in Supplementary Table 7. The target DNA was first chemically synthesized and confirmed by Sanger sequencing and then PCR amplified. The *prfA* sequence was first PCR amplified with forward and reverse amplification primers (NEB Q5® High-Fidelity 2X Master Mix, 95°C 2 min activation, 32 cycles of 94°C denaturing 15 seconds, and 60°C annealing and extension 30 seconds) and then purified with agarose-gel electrophoresis. Recovered DNA concentration was determined by Qubit (ThermoFisher, USA) and later serially diluted while the final concentration was validated by Bio-Rad QX200 ddPCR platform.

For dual-channel assays, template DNA was extracted from human blood samples using QIAGEN DNeasy® Blood & Tissue Kit. See Supplementary Table 1, 5, 7 and 9 for the oligo nucleotides used in the assays.

*2.2 Protocol*

Each CLEAR-dPCR reaction mix is 16 µl in total volume, mixed by 12 µl reaction mix and 4 µl DNA sample. Due to the difference in oligo nucleotide concentration, we slightly modified the betaine concentration for different dPCR assays. The droplet system is very sensitive to surfactant, and we found that Thermo Platinum™ Taq Pol. (Cat. No. 10966026) is compatible with our system.

Simplex dPCR

| | Stock conc. | Vol./ µl | Final conc. in Pre-mix |
|---|---|---|---|
| KCl | 5 M | 0.16 | 50 mM |
| $MgCl_2$ | 1 M | 0.064 | 4 mM |
| Tris-HCl (pH 8.5) | 2 M | 0.16 | 20 mM |
| Forward Primer | 100 µM | 0.16 | 1 µM |
| Reverse Primer | 100 µM | 0.16 | 1 µM |
| Probe | 100 µM | 0.048 | 300 nM |
| dNTP | 10 mM each | 0.64 | 0.4 mM each |
| Betaine | 5 M | 10.08 | 3.15 M |
| Platinum™ Taq Pol. | 1 U/µl | 0.24 | 0.15 U/µl |
| $H_2O$ | | 0.288 | |
| DNA | | 4 | |

Dual-loci dPCR

| | Stock conc. | Vol./ µl | Final conc. in Pre-mix |
|---|---|---|---|
| KCl | 5 M | 0.16 | 50 mM |
| $MgCl_2$ | 1 M | 0.064 | 4 mM |
| Tris-HCl (pH 8.5) | 2 M | 0.16 | 20 mM |
| Forward Primer 1 | 100 µM | 0.136 | 850 nM |
| Reverse Primer 1 | 100 µM | 0.136 | 850 nM |
| Probe 1 | 100 µM | 0.04 | 250 nM |
| Forward Primer 2 | 100 µM | 0.136 | 850 nM |
| Reverse Primer 2 | 100 µM | 0.136 | 850 nM |
| Probe 2 | 100 µM | 0.04 | 250 nM |
| dNTP | 10 mM each | 0.64 | 0.4 mM each |
| Betaine | 5 M | 9.85 | 3.08 M |
| Platinum™ Taq Pol. | 1 U/µl | 0.24 | 0.15 U/µl |
| $H_2O$ | | 0.262 | |
| DNA | | 4 | |

Genotyping dPCR

| | Stock conc. | Vol./ µl | Final conc. in Pre-mix |
|---|---|---|---|
| KCl | 5 M | 0.16 | 50 mM |
| $MgCl_2$ | 1 M | 0.064 | 4 mM |
| Tris-HCl (pH 8.5) | 2 M | 0.16 | 20 mM |
| Forward Primer | 100 µM | 0.144 | 900 nM |
| Reverse Primer | 100 µM | 0.144 | 900 nM |
| Probe 1 | 100 µM | 0.04 | 250 nM |
| Probe 2 | 100 µM | 0.04 | 250 nM |
| dNTP | 10 mM each | 0.64 | 0.4 mM each |
| Betaine | 5 M | 9.92 | 3.10 M |
| Platinum™ Taq Pol. | 1 U/µl | 0.24 | 0.15 U/µl |
| $H_2O$ | | 0.448 | |
| DNA | | 4 | |

Once prepared, the dPCR reaction mixes are immediately loaded into the MiCA droplet generating devices. During the 4-min centrifugation (15,000 rcf), water-in-oil droplets are generated in a 200-μl centrifuge PCR tube (Axygen, PCR02LC) pre-filled with 240 μl receiving silicone oil. Under such centrifuge condition, generated droplets are 41 μm in diameter and we thus estimate that 16-μl dPCR reaction mix has about $4.43 \times 10^5$ droplets. With caps closed, tubes with clear emulsion are then went through PCR thermal cycling: 25°C 2 min for surfactant encapsulation, 95°C 2 min for enzyme activation, 40 cycles (15 s at 92°C and 30 s at 58°C) for amplification.

#### 3. MiCA centrifugal droplet generation free of DNA adsorption

To test if DNA adsorption or loss happens during the centrifugal droplet generation process, we quantified the DNA before and after the centrifugation. We diluted male human genomic DNA into two different concentrations (Sample 1 and Sample 2). From each sample, we took an aliquot (MiCA) and span it through MiCA into an empty tube, while kept another aliquot (NP) untreated. We then quantified the concentrations of the aliquots by Bio-Rad QX200. The assay examined two sites, one on chromosome 1 and the other on chromosome Y (see Supplementary Fig. 1 for detail).

#### 4. The optical setup of readout system

##### *4.1 The light path*

We demonstrate high-throughput, parallel droplet digital PCR readout using a customized light-sheet fluorescence microscope (LSFM, Supplementary Fig. 4). A dual-wavelengths, fiber-coupled semiconductor laser (532/488 nm, OBIS, Coherent®, California) was used as excitation source. The laser was first collimated into a Gaussian beam with diameter ~3.5 mm ($1/e^2$ value). Then a sandwich structure containing an aspherical lens (f = 8 mm) and two cylindrical lenses (CL1 and CL2, f = 20 mm and 12.7 mm, respectively) was designed to transform the round beam into an elliptical shape with a short optical path. The expansion ratio in z and y direction is ×0.4 and ×1.6, respectively, forming an elliptical beam with 5.6 by 1.4 mm in size (Supplementary Fig. 5). A pair of adjustable mechanical slits (0-8 mm aperture size) were placed orthogonally to further truncate the beam and thereby tune the height and thickness of the laser-sheet. The illuminating cylindrical lens (CL3, f = 50 mm) finally produces a large-and-wide laser-sheet that optically sections the droplets layer by layer.

A ×2 infinity-corrected, wide-field detection path (Nikon Plan Apo Fluor 4×/0.13 objective + Thorlabs TTL100 tube lens) was built orthogonal to the plane-illumination path, to collect the fluorescent signals. The centrifuge tube containing droplet samples was mounted onto a motorized translational stage via a tube-holder. During the signal read-out, the tube together with the PCR droplets was rapidly scanned across the laser-sheet along z-axis, typically in a few seconds. A digital camera (Hamamatsu Orca Flash 4.0 v2) continuously recorded the images from the consecutively illuminated planes at a high speed up to several hundreds of frames per second.

*4.2 Components of the readout system*

The whole system can be built upon commercial opto-mechanical components, excepts a few customized parts such as tube holder. Compared to commercial ddPCR instruments, it is significantly more compact (~30×30×10 cm) and cost-effective. We also note that the expense of the system could be further cut down by replacing the current sCMOS camera with an inexpensive one.

*4.3 Characterization of the setup*

The geometry of the laser-sheet, such as its coverage range and axial extension, should be matched to the droplet samples. We thus characterized the laser-sheet properties by imaging a suspension of sub-resolution nano-beads (~500 nm in diameter). The x-z plane of the beads is reconstructed to evaluate the performance of light-sheet. The longitudinal extent of the resolved beads indicating the axial resolution of the system is measured ~22 μm at FWHM value (depth), which is sufficient to resolve the single droplets with ~41 μm diameter (Supplementary Fig. 5a). Furthermore, there is no significant size change of the resolved beads over a wide range of 6.7 mm in x direction (Supplementary Fig. 5a,c), indicating a uniform optical sectioning of the entire droplet stack along the laser propagation direction (width). Using 1 μM fluorescein solution in a rectangular cuvette as target, we then obtain the image of light-sheet illuminated plane to measure the intensity distribution of the laser-sheet along y direction. A stable intensity over 6 mm height is shown to verify a uniform illumination of the laser-sheet along y direction (height), along with an image of a laser-sheet illuminated actual CLEAR dPCR sample (Fig. 3c).

#### 5. 3D image-based droplet counting

We post-processed the image stacks to obtain the accurate copy number of DNA template input for each sample. Due to the slight variation of illumination intensity along vertical direction (y direction), the images were first corrected according to the measured light-sheet intensity profile (Fig. 3c, Supplementary Fig. 6a). Then, we transformed the 3D images in Fourier space to subtract the noise components. The denoised images were obtained by an inverse Fast Fourier transformation (FFT) (Supplementary Fig. 6b). At the identification step, the local maximal-intensity voxels were searched and localized in the 3-D matrices, identified as positive signal candidates (Supplementary Fig. 6c). These located candidates were further processed by an intensity-based thresholding, followed by merging adjacent maximal points (≤ 3 voxels). The final points were fit to spheres for positive identification. The following table shows the staged counting results.

| Stages | In the selected volume | In the whole volume |
|---|---|---|
| Raw images: candidate | 996 | 4,636,199 |
| FFT denoised: candidate | 18 | 14,700 |
| Final for counting: identified | 16 | 12,216 |

#### 6. Results of CLEAR PCR versus that of QX200 ddPCR

Assays carried out on QX200 ddPCR platform contained the same ratio of DNA sample (1/4) in the final reaction mix.

| | Stock conc. | Vol./ µl |
|---|---|---|
| ddPCR Supermix | 2X | 10 |
| Primer & Probe | 100 µM | varied* |
| DNA | | 5 |
| $H_2O$ | | to 20 µl |

*: the final DNA copy number are the same as those in CLEAR dPCR and in 7500-dPCR.

We serially diluted the pre-amplified 280-bp DNA template to five orders of concentration A to E. Along with negative control sample, they were tested by 7500-qPCR, CLEAR dPCR and QX200 ddPCR.

#### 7. Light-sheet illumination at low photobleaching rate supports repetitive CLEAR dPCR imaging

Noninvasive scanning of sealed tube by selective plane illumination mode brings minimal damage to the clear emulsion samples, in terms of both physical structure and fluorescence signal. We verified this through imaging three parallel CLEAR dPCR samples, with each vial being repetitively scanned for 100 times (Supplementary Fig. 12a). The variation of averaged positive signals was plotted versus the cycles, as shown in Supplementary Fig. 12b. Although a gradual signal decline occurred during the repeated scan, sufficient signals were still retained in the last cycle for digital counting. It was calculated that the averaged signal decline per scanning cycle was less than 0.6%, demonstrating the superior performance from sharp, in-focus plane illumination. Furthermore, the samples' internal pattern of stacking was preserved all the time during the 100-cycle scan (Supplementary Fig. 12a).

#### 8. PCR not interfered by high concentration of betaine in CLEAR dPCR emulsion

We added betaine to 5,6-carboxyfluorescein solution, to adjust its refractive index close enough to that of dPCR emulsion. By MiCA centrifugation method, we generated monodisperse emulsions dissolved with and without fluorescein, respectively. By mixing the two emulsions, we successfully made dPCR emulsion analogues (partial droplets contain fluorescein) and scanned them using LSFM (Supplementary Fig. 3a). Fluorescein concentration in the emulsion varied from 0 to 300 nM, and the fluorescence intensity of analogue emulsion droplets increased with higher fluorescein concentration (Supplementary Fig .3b).

In our dPCR experiments, we used hydrolysis probes, containing FAM and quencher, to indicate the reaction process. Before CLEAR dPCR, the PCR mix droplet with 300 nM hydrolysis FAM probe has the fluorescence intensity equivalent to that of 100-nM analogue droplets, indicating the residue fluorescence intensity due to the incomplete quenching. Hence the quenching efficiency $q$ could be obtained ~ 0.66. If all fluorophores are hydrolyzed from the 300 nM probe, it will generate a 300-nM free fluorescein solution and the intensity change will be equivalent to 300 × 0.66 = 200 nM fluorescein. However, after CLEAR dPCR we observed a droplet intensity equivalent to 225 nM fluorescein, indicating the fluorescence intensity elevation of 300-nM probe was equivalent to 200 – 100 = 125 nM fluorescein. We can then calculate the ratio of hydrolyzed probe ($\alpha$) during CLEAR dPCR process is 125 / 200 = 62.5%. To our empirical knowledge, the hydrolyzed ratio of such probes in qPCR ranges from 40% to 70%, depending on the probe sequence and fluorophore/quencher configuration. Therefore, our experimental data suggest that the polymerase reaction is not significantly interfered by the betaine added into the droplets for RI matching.

#### 9. Lossless dPCR reduces quantitative uncertainty

Conventional quantitative measurements' uncertainty is usually resulted from instrumental errors and their measurement output often comes in a continuous format. However, dPCR quantification has been accurate enough for us to care for errors that occurs unconventionally and those long been neglected, subsampling errors. Besides partition volume variation and insensitivity or lack of detection resolution in signal acquisition, the randomness in the subsampling and partitioning processes in dPCR are also sources of uncertainty, which is discussed hereinafter.

The probability of having $k$ molecules in the sample follows a Poisson distribution,

$$\Pr(k) = \mu^k \frac{e^{-\mu}}{k!}$$

Where $\mu$ is the expected copy number per each draw/reaction. The actual number of molecules drawn tends to be centered to the expectance as the concentration increases (Supplementary Fig. 7a), and the uncertainty introduced in subsampling is dependent upon sample concentration. Subsampling error is prevalent in almost every measurement as long as sampling is done to only part of the target of interest rather than to the entirety.

Another source of uncertainty is the partitioning process, which is also a random process and may result in different positive partition numbers even the number of molecules drawn in each reaction is the same. Consider we have $N$ partitions in a reaction, each partition has a volume $v$. The expected number of molecules in the sample $m$ in volume $V$ are,

$$V = Nv$$
$$m = CV$$

Where $C$ is the volumetric concentration of target molecule. It is noted that due to the aforementioned subsampling error, $m$ and $C$ are random variables themselves. Suppose $P$ partitions are found positive after amplification (no partition loss), and we define two parameters

$$\lambda = {}^{m}/_{N} \text{ and } p = {}^{P}/_{N}$$

Because $\lambda = Cv$, the expectance of the number of molecules in each partition is $\lambda$. And the likelihood of a given partition has at least one molecule is

$$\sum_{k \neq 0} \Pr = 1 - \Pr(0) = 1 - e^{-\lambda}$$

Looking at the partitions as whole, for $N$ partitions investigated here, the number of positive partitions $P$ follows binomial distribution $\mathrm{B}(N, 1 - e^{-\lambda})$, whose expectance $\mathrm{E}(P)$ and standard deviation $\sigma_P$ are

$$\mathrm{E}(P) = (1 - e^{-\lambda})N$$

$$\sigma_P = \sqrt{e^{-\lambda}(1-e^{-\lambda})N}$$

And because of the definition that $p = {}^{P}/_{N}$,

$$\hat{p} = \hat{P}/\mathrm{N}$$

$$\sigma_p = \sqrt{(1-p)p/N}$$

In practice, $P$ is the direct measurand we first obtain and $p$ after that, and we want $\lambda$ for calculation of the concentration. We have the confidence range for $p$,

$$[p - z_\alpha\sqrt{\frac{(1-p)p}{N}}, p + z_\alpha\sqrt{\frac{(1-p)p}{N}}]$$

And because $\mathrm{E}(P) = (1-e^{-\lambda})N$,

$$\hat{\lambda} = -\ln(1-p)$$

When confidence level $\alpha = 5\%$, $z_\alpha$=1.96. Plug the boundary values of $p$ into the equation $\hat{\lambda} = -\ln(1-p)$ and we have the confidence of $\lambda$. The uncertainty of a measurement is usually defined as,

$$U = \frac{\lambda_{\text{upper}} - \lambda_{\text{lower}}}{\lambda}$$

The smoothed curves in Supplementary Fig. 7b shows the uncertainty level of dPCR assays with respect to varied molecule volumetric concentration, and with different effective partition number, obtained as theoretical values calculated using the equation above. While the dots in Supplementary Fig. 7b are the results from a Monte-Carlo simulation on Matlab. For each assay (each volumetric concentration point, each partition number point) we simulated 10,000 repeats of 'dPCR assay' by generating binomial random variables. After sorting the results of the repeats, we use the average value of $\lambda$ in the 250$^{th}$ and 251$^{st}$ places as lower boundary, and that of 9750$^{th}$ and 9751$^{st}$ as upper boundary. It is clear that the Monte-Carlo simulated uncertainty level is in good accordance with theoretical results, which validated our Monte-Carlo model as useful tool to estimate dPCR uncertainty. This Monte-Carlo model was then used in the following discussions on dPCR WITH partition loss.

We adjusted the Monte-Carlo model to only count only a portion of the partitions in order for the uncertainty levels in dPCR with partition loss. As shown in Supplementary Fig. 7c, the loss rate directly boosts uncertainty throughout the input range of measurand concentration. It is clear that in lossless cases, as the molecule concentration increases, assays with less partition numbers are less capable of quantifying sample with high concentration (Supplementary Fig. 7c). Numbered hollow dots respectively correspond to the concentrations in the absolute quantification assay in Fig. 4b,c to show their theoretical values of uncertainty at this concentration. Although CLEAR dPCR has a loss rate = 0, we still drew the curves with loss rate above that value in order to show how loss of droplet influences the uncertainty of the measurement. Due to the inevitable existence of droplet loss, the uncertainty level of QX200 is higher than lossless CLEAR dPCR throughout the measurand concentration. And the larger partition number further extends effective concentration range of dPCR to more than 10-fold increase.
